# Losses outlast gains in Northern Hemisphere tree growth under climate change

**DOI:** 10.64898/2026.09.06.749676

**Authors:** Erez Feuer, Stefan Klesse, Guy Amir, Nadav Peleg, Yair Mau

**Affiliations:** Institute of Environmental Sciences, The Hebrew University of Jerusalem, Rehovot, Israel; Swiss Federal Institute for Forest, Snow and Landscape Research WSL, Birmensdorf, Switzerland; Department of Computer Science, Cornell University, Ithaca, NY, USA; Institute of Earth Surface Dynamics, University of Lausanne, Lausanne, Switzerland; Expertise Center for Climate Extremes, University of Lausanne, Lausanne, Switzerland

## Abstract

Under climate change, will forests grow more or less? The answer differs from place to place, obscuring the overall direction. Here we trace it by learning nonlinear climate–tree-growth relationships from 4,110 Northern Hemisphere tree-ring chronologies recorded since 1902, spanning more than 12 million ring-width measurements, and projecting them through the twenty-first century under CMIP6 scenarios. Projected growth declines, and gains and losses are temporally asymmetric: once losses emerge, they are rarely reversed (90% retained), whereas gains fade (56% retained). This asymmetry holds in every well-sampled region and strengthens under higher forcing. The decline itself is driven by warming and rising atmospheric water demand, whose broadly negative contribution outweighs precipitation-driven gains. Across the Northern Hemisphere, growth does not just move between gains and losses; it is drawn steadily downward.

---

Forests are a major component of the terrestrial carbon cycle and influence climate through both carbon storage and biophysical feedbacks (6, 43). In turn, climate change reshapes the thermal and hydrological conditions under which forests grow, shifting both average climate and extremes, with consequences for growth, physiological function and survival (1, 36, 8). These outcomes are neither uniformly positive nor negative; thus, response of tree growth to climate change remains difficult to generalize.

This difficulty arises because climate–tree-growth relationships typically follow nonlinear response curves (10, 15, 4, 53). The impact of climate change — the change in growth it produces — depends not only on the magnitude of a climate shift but also on where along the response curve that shift occurs, so the same change can produce positive, negative, large, or small impacts (Fig. 1a). In multidimensional climate space, this starting point includes the state of other climate variables: a change in one dimension can have different impacts depending on the background climate (Fig. 1b). Together, these relationships define a context-dependent response surface across which progressive climate shifts translate into impacts on growth that may change in magnitude or direction over time (Fig. 1c).

**Figure 1:**
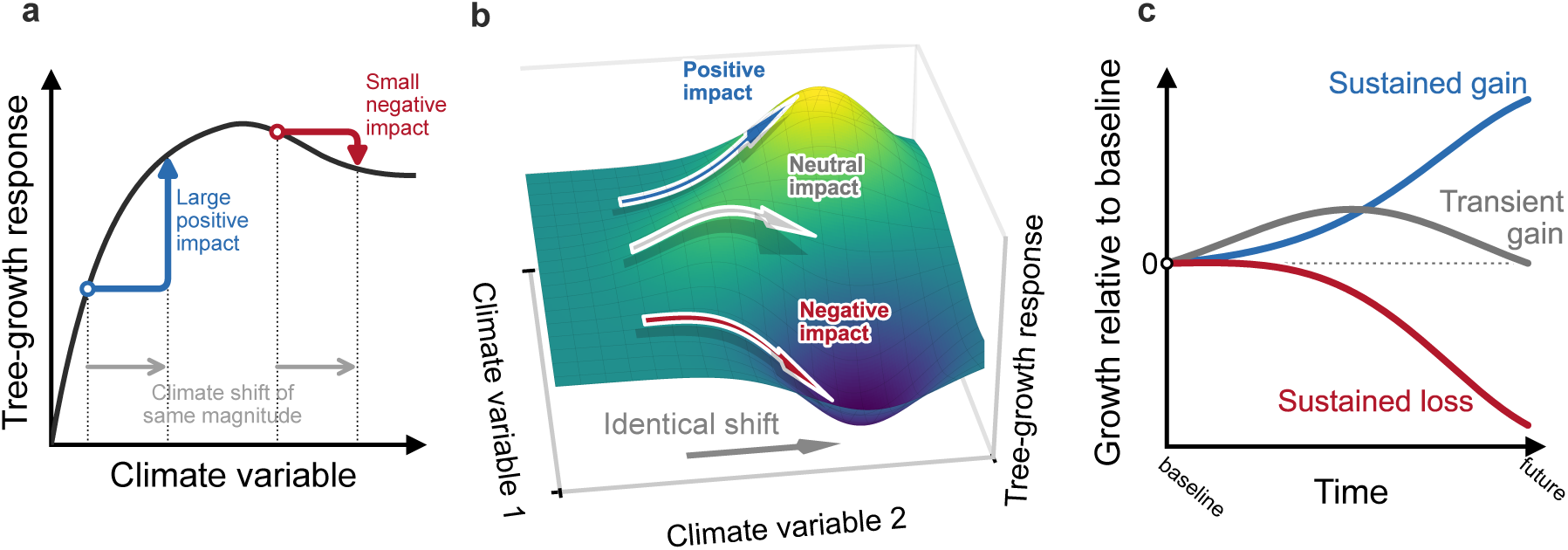
Nonlinear climate–tree-growth responses shape divergent growth trajectories under identical climate shifts. Schematic of the conceptual framework linking nonlinear climate–tree-growth relationships to the growth impact of climate change. (a) The impact of a climate shift — the change in predicted growth it produces — depends on where it occurs along a nonlinear response curve: the same shift can raise growth on a rising portion of the curve or lower it beyond an optimum. (b) In multidimensional climate space, impact also depends on the background state of other climate variables, so an identical shift in one dimension can produce positive, neutral, or negative responses under different background conditions. (c) Viewed through time as growth relative to baseline, the same response geometry yields different trajectories: sustained gains, transient gains that return toward baseline, or sustained losses.

How these impacts evolve depends on the balance among the three principal climatic controls on tree growth: air temperature, precipitation, and atmospheric water demand (51, 4, 20, 19). Where low temperature or a short growing season limits growth, warming can enhance it; but as warming continues, rising atmospheric water demand can carry the same forests from this energy-limited regime toward water limitation, where further warming suppresses growth (15, 4, 50, 49). Precipitation affects growth through its contribution to water supply, but realized water availability depends on both supply and atmospheric demand; when this balance shifts toward deficit, it can suppress growth (25, 56). Rising atmospheric water demand, expressed as increasing vapor pressure deficit (VPD), can suppress growth by driving water loss and stomatal closure, with the strongest effects during droughts (40, 20, 55, 38). Because climate change shifts these three controls simultaneously, the open question is how their effects combine, whether they offset, amplify, or reverse one another, to determine the overall growth response.

Global projections of vegetation response to climate change already exist, but they largely target carbon uptake or canopy greenness rather than directly measured tree growth, from which they are often decoupled (9, 18). For growth itself, as captured by tree-ring measurements, the literature has largely followed two paths. Regional and continental studies have projected growth under climate scenarios, in some cases capturing nonlinear responses, but resolving these relationships in detail only for particular regions or species (31, 15, 10, 28, 29, 35). Broad cross–continent tree-ring syntheses have described growth sensitivity to climate, focusing on linear relationships within historical climate (4, 56). These lines of work have stayed apart: where coverage has been broad, the response has been treated linearly or left unprojected; where it has been modeled nonlinearly or projected forward, the scope has remained regional or continental. Our aim here is to bring them together, i.e., to characterize nonlinear climate–tree-growth response surfaces across a broad, multi-biome tree-ring network, and to project how twenty-first-century climate change moves forests across them.

We compiled a broad, multi-biome dataset of 4,110 Northern Hemisphere ring-width index (RWI) chronologies, drawn primarily from the International Tree-Ring Data Bank (ITRDB) (39) with additional European beech records (29), representing more than 12 million tree-ring observations. As climatic predictors, we used precipitation, maximum temperature, and VPD from the observed CRU TS v4.09 record (21), each represented by four 3-month summaries over the annual window most influential for ring formation (44). We estimated empirical climate–tree-growth response surfaces with XGBoost, fitting RWI against these predictors for 1902–1980 and allowing nonlinearities and interactions to be learned from the observations rather than prescribed. We then applied the model to observed climate and CMIP6 projections under SSP2-4.5, with the high-forcing SSP5-8.5 as a limiting case, estimating growth impacts as changes in predicted RWI relative to the 1902–1980 baseline.

Because our response variable is detrended RWI, the model diagnoses climate-driven changes in radial growth rather than absolute wood production or carbon uptake (3, 9). We find that projected climate change decreases growth overall, with losses outlasting gains across regions through the twenty-first century. Decomposing these responses shows that the decline is driven primarily by a broadly negative warming–demand contribution that often outweighs positive water-supply contributions.

## Projected growth declines and broadens

To evaluate the combined impact of change across all climate predictors, we applied the model to 30-year climate windows extending from observed CRU TS climate into CMIP6 projections. Throughout, we summarize responses on a 1^◦^ grid, assigning each chronology to a grid cell and taking the median response within it, so that densely sampled regions do not dominate (Methods). Relative to the baseline, predicted RWI declined progressively through the twenty-first century (Fig. 2a). As projected climate change intensified, with binned-grid maximum-temperature anomalies reaching +3.7 ^◦^C, statistically significant growth changes emerged predominantly below baseline: by late century (2071–2100), 37% of grid cells declined against 10% that gained (Fig. 2b).

**Figure 2:**
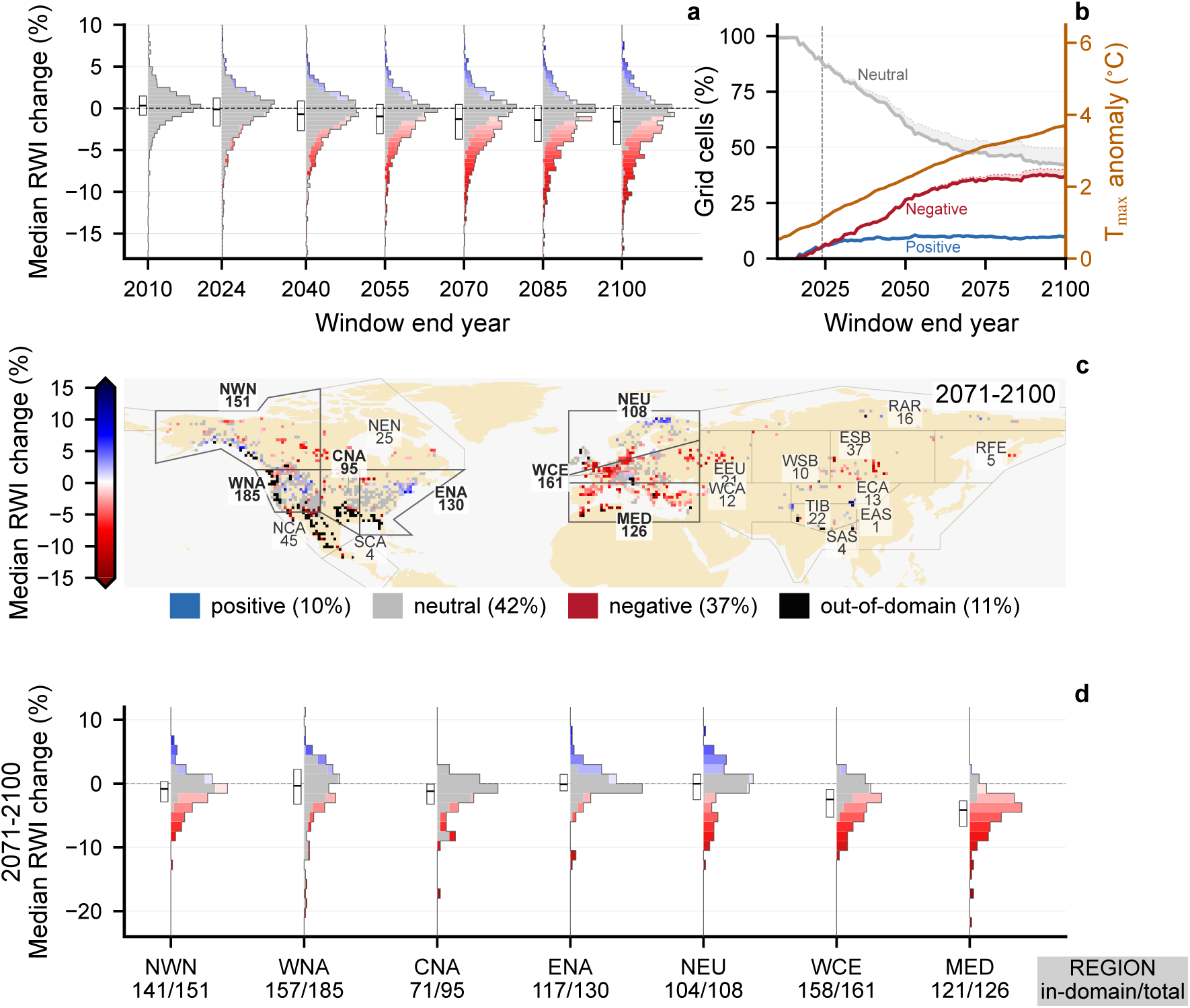
Projected radial-growth responses decline and broaden asymmetrically under future climate change. (a) Distributions of one-degree binned grid-cell median predicted relative ring-width index (RWI) change relative to the 1902–1980 baseline, calculated in rolling 30-year windows from 1981–2010 through 2071–2100 using observed CRU TS climate followed by CMIP6 SSP2-4.5 projections. Predicted RWI change is the percentage change in a chronol-ogy’s XGBoost-predicted ring-width index in a target window relative to its 1902–1980 baseline (% ΔRWI), aggregated to grid cells; a value of −4% means growth 4% below baseline. For projected windows, grid-cell change is the median across the twelve CMIP6 models of each model’s 30-year median. Colors follow the map scale and indicate predicted RWI change; gray histogram components are in-domain cells whose change was not statistically significant after Benjamini–Hochberg FDR correction (*q >* 0.05), and colored components are statistically significant cells (*q* ≤ 0.05). Boxes show the in-domain interquartile range and median for each window. (b) Rolling-window fractions of mapped grid cells with negative, neutral, or positive responses, with the binned-grid annual maximum-temperature anomaly relative to baseline on the right-hand axis. For each fraction, the shaded band extends the solid in-domain line to include cells frozen at their last in-domain state after leaving the climate domain. (c) Late-century map of binned median predicted RWI change. Colored cells are in-domain grid cells with statistically significant change; gray cells are in-domain but not significant, and black cells are outside the training climate domain. Percentages indicate the share of mapped grid cells with positive change, neutral (not significant) change, negative change, and out-of-domain climate. AR6 reference-region boundaries are overlaid, with darker boundaries marking the well-sampled regions summarized in panel d. (d) Late-century in-domain grid-cell response distributions for AR6 reference regions with at least 95 mapped grid cells. Histograms use the same significance encoding as panel a, with gray components for in-domain cells that are not significant and colored components for statistically significant cells. Boxes show each region’s in-domain interquartile range and median. Grid-cell binning reduces over-representation of highly sampled regions in the tree-ring network.

The distribution in RWI change both decreased and widened, with its interquartile range increasing from 2.36 to 4.75 percentage points. This widening was asymmetric: both the lower and upper quartiles shifted toward greater losses, but the lower moved roughly three times farther than the upper (3.54 versus 1.15 percentage points). Under the high-forcing SSP5-8.5 scenario, these changes were amplified: the median declined further, and the upper quartile shifted below zero (Fig. S1a). The widening was also evident within individual CMIP6 models, rather than arising from intermodel disagreement (Fig. S2).

These late-century responses were spatially structured (Fig. 2c,d). We summarized regional responses for the seven IPCC AR6 reference regions with at least 95 mapped cells (24) (Fig. 2d). Decline was strongest and most consistent in the Mediterranean (MED) and in western and central Europe (WCE), where statistically significant negative responses made up 85% and 54% of mapped cells and median growth fell by 4.2% and 2.5%. Other regions showed weaker or mixed responses: in eastern North America (ENA) and northern Europe (NEU), significant positive and negative responses were comparably frequent. In several regions, including central North America (CNA), the change in most cells was not significant, with projected responses remaining within baseline variability. In other regions, most notably northern Central America (NCA), cells entered out-of-domain (OOD) climate space not represented during model training and were excluded to avoid unsupported extrapolation (see Methods); OOD conditions became considerably more widespread under SSP5-8.5 (Figs. S1b and S4).

Across the network, then, projected growth declined overall but unevenly: decisively in parts of Europe, weakly or mixed elsewhere, and within baseline variability across much of interior North America. These distributions, however, count cells at each point in time without following what becomes of any individual cell, and cannot show whether a given departure from baseline endures or reverses. We therefore tracked how each cell evolved from the baseline through the end of the century.

## Losses are retained while gains fade

To follow individual grid-cell trajectories, we tracked them from their first departure into a positive or negative state to the end of the century, retaining the last in-domain state for cells that subsequently left the training climate domain (Fig. 3a). Negative states rarely reverted: 90% remained negative by 2100 (95% CI 87–92%), against 56% (95% CI 49–62%) of positive states (Fig. 3b). Under high-forcing SSP5-8.5, the split sharpened further: negative retention rose to 97%, so that almost no decline reversed, while positive retention fell to 46% (Fig. S5b). Negative retention exceeded positive retention in all seven well-sampled AR6 regions (80–100% versus 0–77%; sign test *P* = 0.016), and a region-blocked bootstrap placed the full-network gap at 34 percentage points (95% CI 20–52).

**Figure 3:**
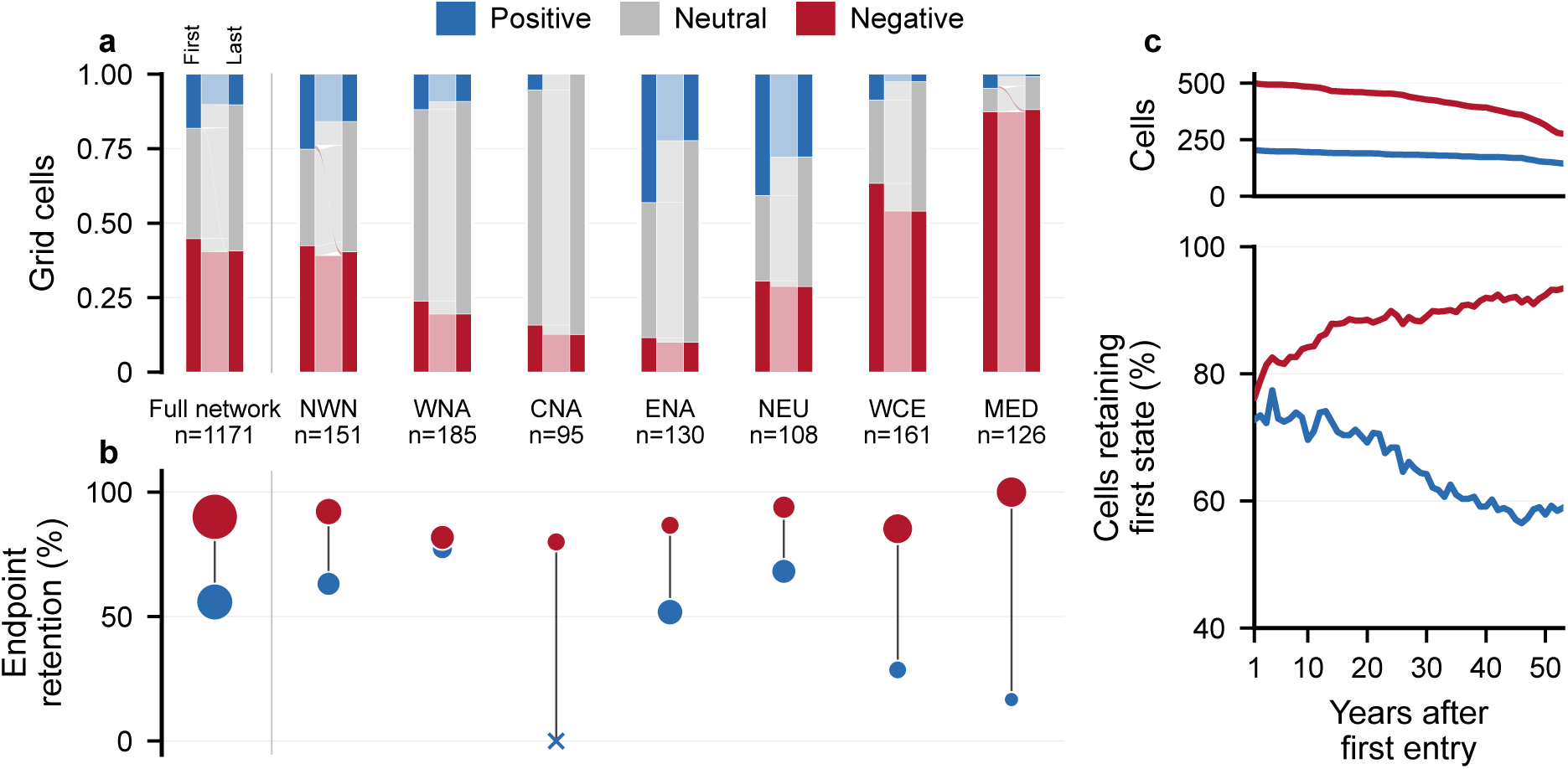
Negative growth responses are retained while gains fade under SSP2-4.5. Each grid cell is tracked from its first departure into a positive or negative state to its last in-domain state, following the rolling 30-year windows; cells that later leave the climate domain are held at that last state (Methods). A cell’s state is negative or positive where its predicted RWI change relative to the 1902–1980 baseline is statistically significant and in-domain, and neutral otherwise. (a) Cell-level transitions from the first non-neutral state to the last state, for the full network and the seven well-sampled AR6 reference regions. Bar heights show the fraction of grid cells in each state at first entry and at the last state. (b) Retention of first-entry positive and negative states: the fraction of cells still on the same side of the boundary at their last state, with point size scaled by the number of first-entry cells. (c) Retention as a function of time since first entry: the fraction of cells still on their first-entry side, with each cell aligned to its own first entry rather than to a common year, for positive (blue) and negative (red) entries. At each horizon, a cell is counted only while it remains observable and in-domain; cells drop out as they leave the training climate domain or reach the end of the century, and curves are shown only where at least 150 cells of each sign remain (Methods). The upper strip shows the number of observable cells of each sign at each horizon, which declines as cells drop out. Negative retention rises and positive retention falls as time since entry lengthens, widening the gap between them.

This asymmetry arose from retention rather than entry: in eastern North America (ENA) and northern Europe (NEU), cells entered positive states more often than negative ones (43% versus 12% and 41% versus 31%, respectively), yet negative states were retained more often through 2100 (Fig. 3b). Following cells by time since their first entry rather than to a fixed date, the two sides diverged: negative states grew more persistent while positive states decayed, the gap widening steadily with time since entry (Fig. 3c). By 50 years after entry, 92% of negative states were retained against 58% of positive states under SSP2-4.5, and 97% against 47% under SSP5-8.5 (Fig. S5c). Losses therefore consolidate while gains fade with time since departure.

When either state was left, cells almost always returned to neutral rather than crossing to the opposite sign. Under SSP2-4.5, positive cells returned to neutral in 43% of cases and negative cells in 10%, while direct crossings from one sign to the other were rare in both directions (1% positive-to-negative and under 1% in the reverse direction) (Fig. 3a). This directional imbalance strengthened under high-forcing SSP5-8.5, where downward crossings rose to 14% and upward crossings fell to zero (Fig. S5a).

These unequal transition dynamics caused negative responses to accumulate while positive responses dissipated. We next trace these changes to their climatic origin.

## Warming and VPD drive the losses

We decomposed projected changes into a water-supply contribution from precipitation and a warming–demand contribution from maximum temperature and VPD, and summarized them at the grid-cell level. Because maximum temperature and VPD are strongly correlated, their individual contributions could not be resolved reliably and were therefore grouped. For the end of the century (2071–2100), both water-supply and warming–demand contributions were expressed relative to baseline as changes in SHAP (Shapley additive explanations; see Methods) values (ΔSHAP; Fig. 4a). The two contributions were opposite in sign across most of the network: under SSP2-4.5, warming–demand was negative in 91% of grid cells while water-supply was positive in 79%.

**Figure 4:**
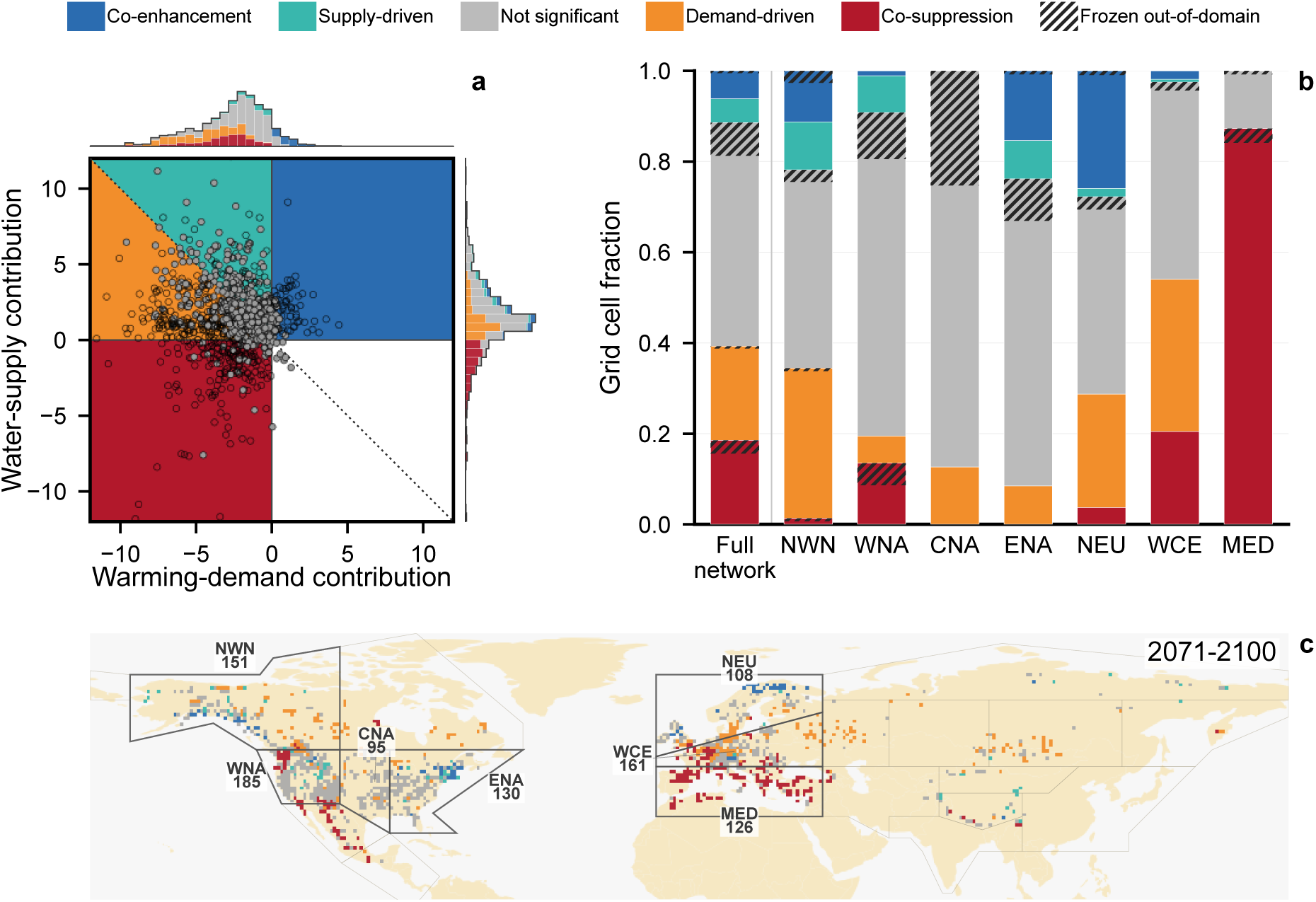
A negative warming–demand contribution drives the projected decline against a positive water-supply contribution. For each grid cell, we ask which climate drivers produced its projected growth change: how much of the change is attributable to water supply (precipitation) versus warming–demand (maximum temperature and VPD). This driver attribution is the change in each group’s SHAP contribution from the baseline (ΔSHAP), expressed in the same percent-equivalent RWI units as the growth change itself, so a driver’s contribution reads directly as the growth it adds or removes. These contributions are additive on the underlying log-RWI scale; after conversion to percent-equivalent units they are interpretable individually but do not sum exactly (Methods). Contributions are taken as the median across models and summarized within 1-degree grid cells. Cells are then assigned to five regimes from the signs of the two contributions and the climate net: co-enhancement and co-suppression (both contributions of the same sign), supply-driven and demand-driven (opposing contributions with a positive or negative climate net), and neutral (in-domain but not statistically significant). (a) Late-century (2071–2100) contribution plane under SSP2-4.5. Points show in-domain grid cells; statistically significant non-neutral cells are black and neutral cells gray. Background colors show the regime partitions, and marginal histograms show the regime composition along each contribution axis. (b) Late-century regime composition for the full sampled network and the seven well-sampled AR6 reference regions. Bars show the fraction of grid cells in each regime; hatching marks cells held at their last in-domain regime after leaving the climate domain. (c) Geographic distribution of late-century regimes, shown as the modal regime within each 1-degree grid cell, with AR6 reference-region boundaries overlaid.

We partitioned the contribution plane into five regimes: co-enhancement and co-suppression, where both contributions acted in the same direction; supply- and demand-driven responses, where opposing contributions yielded a positive or negative climate net; and neutral responses, where the change was not significant. Under SSP2-4.5, demand-driven loss was the most common non-neutral regime across the network, at 243 of 594 non-neutral grid cells (41%), and the leading negative regime in five of the seven well-sampled AR6 regions (Fig. 4b,c). Co-suppression dominated losses only in western North America (WNA) and the Mediterranean (MED), accounting for 87% of cells in the latter.

The balance of contributions within positive responses shifted with forcing, even though positive responses remained a similar fraction of the network (11% under SSP2-4.5 versus 10% under SSP5-8.5). Within these positive responses, the supply-driven fraction rose from 46% under SSP2-4.5 to 61% under SSP5-8.5, at the expense of co-enhancement, though a co-enhanced core persisted even under high forcing (Fig. S6b). Under stronger forcing, the surviving gains depended increasingly on water supply alone, offsetting a warming–demand contribution that had itself turned negative.

The decomposition identifies warming–demand as the broadly negative contribution underlying the projected decline, but not why the same climate change drives some responses up and others down. This divergence is clearest at the level of individual chronologies: it depends on where each chronology begins on the response surface and how projected change displaces it, which we examine next.

## Nonlinearities explain divergent impacts

To determine why projected climate change produced opposing responses among chronologies, we followed their baseline-to-late-century displacements across the learned climate–tree-growth surface. Four predictors (P_4–6_, D_4–6_, T_7–9_, and T_10–12_) accounted for approximately 53% of the summed absolute change in attribution among climate predictors by late century (Fig. S7). Their pairwise response surfaces, precipitation–VPD and VPD–temperature, therefore capture much of the geometry underlying the divergence (Fig. 5).

**Figure 5:**
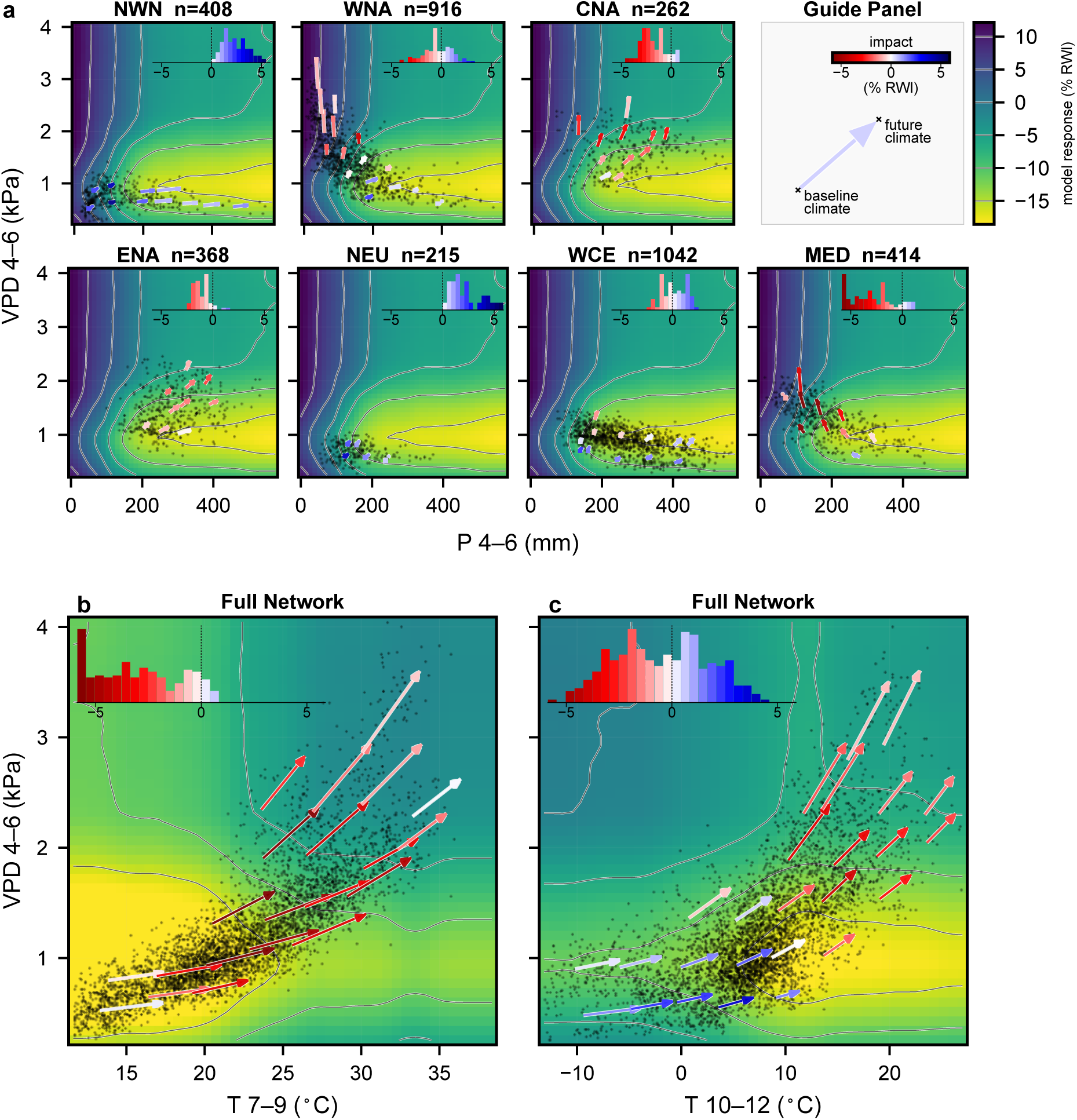
Projected growth impact depends on how climate change moves chronologies across the nonlinear surface. Each panel is the model’s learned growth response over a pair of climate predictors. Arrows indicate how groups of chronologies move through climate space, from their mean present-day climate (base) to their mean late-century climate (tip). Arrow color denotes the impact on growth, with red shades indicating declines and blue shades increases. The background surface sets the terrain the arrows cross (see Methods), with gray contour lines marking levels of equal predicted growth. The impact of a climate shift depends on how the arrows meet these contours: an arrow crossing them changes growth sharply, while one running parallel to them barely changes it. Black dots mark the observed baseline climates of the underlying chronologies and show where the surface is supported by data; the surface should not be read where no dots fall. Histograms inside each panel show the distribution of impacts. (a) April–June VPD (D_4–6_) versus April–June precipitation (P_4–6_), shown separately for the seven well-sampled regions. (b, c) April–June VPD (D_4–6_) versus July–September (T_7–9_) and October–December (T_10–12_) maximum temperature, pooled across the full network. All panels show SSP2-4.5 late-century climate (2071–2100) relative to the 1902–1980 baseline. The full set of pairwise surfaces, and regional breakdowns of the leading pairs, are shown in Figs. S14–S24.

The regional precipitation–VPD surfaces present how contrasting outcomes emerge from the joint climate displacement rather than from either variable alone (Fig. 5a). In northwestern North America (NWN) and northern Europe (NEU), chronologies occupied the lower end of the VPD range, where projected increases in both precipitation and VPD moved them toward higher growth. In central North America (CNA) and eastern North America (ENA), precipitation also increased, but the accompanying rise in VPD carried chronologies across declining portions of the surface and outweighed the positive water-supply contribution. Western North America (WNA) and western and central Europe (WCE) spanned both rising and declining portions of the surface and consequently showed mixed responses. Mediterranean (MED) chronologies combined declining precipitation with increasing VPD, so both components of the displacement acted toward lower growth.

These regional contrasts demonstrate on the learned surface why local climate sensitivity is not equivalent to realized climate-change impact (Figs. 1 and 5a). Sensitivity is the steepness of the surface at a chronology’s current climate. Impact is the growth change actually realized, and it depends not only on that steepness but on the direction and distance climate change carries the chronology across the surface. Western North America (WNA) makes the distinction clear: there, short displacements that cut straight across the response contours changed growth more than longer ones that ran nearly along them (Fig. 5a). Projected impact therefore depends on both displacement magnitude and its alignment with the local response gradient, and cannot be inferred from sensitivity alone.

The same dependence on starting climate and displacement direction structured the response to warming. Displacements across the D_4–6_×T_7–9_ surface were predominantly associated with lower growth, with positive responses confined largely to its coldest and lowest-VPD edge (Fig. 5b). The D_4–6_×T_10–12_ surface instead contained a broader rising region in colder, low-VPD climate space, where warming could increase growth before reaching a crest beyond which further warming and atmospheric demand reduced it (Fig. 5c). The growth effect of warming therefore depended on both the three-month window and the climate state from which the displacement began.

The bounded rising region of the D_4–6_×T_10–12_ surface also provides a mechanism for the lower retention of gains. Chronologies beginning on its rising side can initially be displaced toward higher growth, but continued movement along the same broad climate trajectory carries them toward and eventually beyond the crest. The same progression can therefore produce an early gain followed by declining growth as warming and atmospheric demand intensify. Nonlinear response geometry thus links the concentration of gains in colder, low-demand climate space to their lower persistence under continued change.

Together, these two-dimensional surfaces show that projected climate impact emerges from the interaction between starting climate, the direction and magnitude of climate displacement, and the nonlinear response geometry crossed along the way. Within the learned climate–tree-growth relationships, this geometry shows how positive water-supply contributions can be outweighed by warming–demand losses, and how gains can emerge along part of a climate trajectory but weaken as displacement continues.

## Discussion

Across a broad, multi-biome tree-ring network, twenty-first-century climate change is projected to decrease radial growth, but the central result is a temporal asymmetry: losses, once established, are rarely reversed, whereas gains fade. This asymmetry held in every well-sampled region in the Northern Hemisphere and strengthened under higher forcing. The combined effect of warming and rising atmospheric water demand was negative across most of the network and, in many regions, outweighed precipitation-driven gains, producing widespread losses.

These projections rest on an inherently non-mechanistic partial model: beyond climate, radial growth carries biological trends, disturbance, and other influences a climate model cannot capture (12). Our static predictors (species, elevation, and chronology identity) absorb non-climatic variation, so that the learned climate response need not account for it. A grouped permutation test confirms that this signal is climatic: when the climate–tree-growth link is broken with static structure left intact, model skill collapses to near zero (Methods; Fig. S9). The model learns this response directly from the data with gradient-boosted trees rather than through prescribed functional forms. Across many species and chronologies, it reproduces the expected redistribution of growth limitation from temperature toward water and atmospheric demand (4, 20, 55, 41). The projected changes are numerically small, a few percent of each chronology’s baseline growth, but because ring width is detrended, they are departures from each chronology’s own expected growth, not fractions of a larger total, and they recur as displacement continues. The model underestimates the amplitude of year-to-year climate responses, predicting within-chronology good-minus-bad-year contrasts about two-thirds as large as observed (Methods).

The provisional nature of gains has a structural origin. Regional studies have long reported warming-driven gains that later weaken or reverse in cold-limited and high-latitude forests (15, 50, 35, 49): warming relaxes a temperature limitation and growth rises, but only within a bounded region of climate space, beyond which continued displacement carries a chronology over the peak of its response into decline. Our network resolves these regional findings as one nonlinear response, gain and loss depending on where continued warming carries each chronology relative to the peak. Our decomposition finds other gains resting instead on water supply, sustained while the warming–demand contribution is already negative, consistent with observations of growth held up by precipitation where warming is adverse (14, 34). Across both, projected gains are provisional — sustained only where a favorable balance holds, and eroded as rising warming and atmospheric demand tip it.

Several boundaries constrain this interpretation. Detrended ring width records climate-driven changes in radial growth, not absolute wood production or carbon uptake, from which drought and allocation shifts can decouple it (9, 26, 7); it does, however, track the increment of above-ground woody biomass, the pool in which trees hold most of their carbon (2, 45). Our projections depend on the climate models and scenarios that drive them; we use twelve CMIP6 models spanning different families, which sample structural climate-model uncertainty, and headline the mid-range SSP2-4.5, closest to current emission trajectories (22), with high-forcing SSP5-8.5 as an upper bound. A deeper assumption is space-for-time substitution: as a chronology’s climate changes, it moves into conditions the model has learned from other sites that experience them today (28, 17). Because ring width is standardized, the model uses relative growth within each chronology rather than absolute differences between chronologies. We hold species, elevation, and growth history fixed, project relative to each chronology’s own baseline, and, because a data-driven model cannot reliably extrapolate beyond its training climate, withhold prediction beyond the training climate domain — the range of climate the model was trained on. Our network is geographically and taxonomically uneven and biased toward climate-sensitive sites (27, 3), with sparse coverage across much of the globe, so inference is strongest where sampling is densest. Finally, several processes lie outside the model altogether: we do not simulate CO_2_ fertilization, nor biotic and abiotic disturbances (such as fire and insect outbreaks), mortality, or species turnover (46, 1, 36), and our three-month predictors represent sub-seasonal extremes only indirectly, so brief events such as heatwaves, rainfall pulses and compound episodes are captured only in part.

Here we bring together three strengths that had developed apart: resolving climate–tree-growth responses in their nonlinear detail, measuring them across a broad, hemisphere-wide network, and projecting them into the future. Together they reveal that the growth of Northern Hemisphere trees is reduced more often than enhanced, and once reduced, it tends to persist. We looked here only at how growth responds to shifts in mean climate; how that combines with the many processes we set aside, from CO_2_ fertilization to the disturbances, mortality and extremes escalating under climate change, is a promising direction for future work.

## Methods

### Tree-ring data

Tree-ring width measurements were obtained primarily from the NOAA International Tree-Ring Data Bank (ITRDB) (39), supplemented by a European beech network (29). Within each source record, measurement series shorter than 20 years were excluded, and the remaining series were screened for crossdating consistency, detrended with a cubic smoothing spline using dplpy (42), and averaged with a biweight robust mean into standard ring-width index (RWI) chronologies. The spline had a 50% frequency-response cutoff at 50 years (13), so it removes the low-frequency growth trend while preserving the interannual variability on which the climate–tree-growth model is fit; because some low-frequency variability is climatically forced, this detrending may make the magnitude of the projected growth responses conservative while leaving the interannual response intact. Chronology years were retained from 1901 onward where sample depth was at least 10 series. After matching to climate data and applying the model-fitting filters described below, the final network comprised 4,110 Northern Hemisphere RWI chronologies from 70 species, representing more than 12 million ring-width measurements. Retained chronology-year observations had a median sample depth of 27 series (interquartile range, 19–39).

### Climate data and projections

Observed climate was derived from CRU TS v4.09 monthly fields for 1901–2024 (21). Chronology coordinates were assigned to the nearest 0.5^◦^ CRU grid cell. From CRU TS, we extracted monthly mean daily maximum temperature (^◦^C), total monthly precipitation (mm month^−1^) and monthly vapor pressure (hPa). We derived monthly VPD (kPa) from monthly mean daily maximum temperature and monthly vapor pressure (see Supplementary Section S1). Monthly values were aggregated into four non-overlapping three-month windows: January–March (months 1–3), April–June (4–6), July–September (7–9) and October–December (10–12). Maximum temperature and VPD were averaged within each window, whereas precipitation was summed.

Future climate data were obtained from 12 general circulation models (GCMs) in the NEX-GDDP-CMIP6 dataset under SSP2-4.5 and SSP5-8.5 (48, 47). Daily values were extracted at the nearest 0.25^◦^ NEX-GDDP-CMIP6 grid cell for each chronology and aggregated to monthly site series. Projected VPD was calculated from daily maximum temperature and relative humidity before monthly aggregation. We then bias-adjusted each model’s monthly series to the CRU reference over 1950–2014 (23), separately for each model, site, calendar month and variable: additive monthly offsets for temperature and VPD, and multiplicative monthly factors for precipitation. Corrected monthly projections were aggregated to the same three-month predictors as the observed climate. The CMIP6 ensemble is listed in Supplementary Table S1.

### Predictors and model fitting

We trained an XGBoost regression model (11) to predict annual log-transformed RWI from three-month climate summaries, chronology context, and previous-year growth. The response variable was

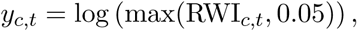

where the small floor of 0.05 prevents undefined values for zero RWI observations. The log transformation makes proportional changes symmetric around RWI = 1, so that halving and doubling RWI have equal magnitude and opposite signs. The fitted model can be written as

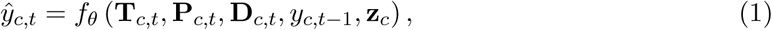

where *c* indexes chronologies and *t* the growth year. The vectors **T**, **P** and **D** contain three-month summaries of maximum temperature, precipitation and VPD, *y_c,t_*_−1_ is the previous-year log RWI, and **z***_c_* holds chronology identity, species identity and elevation. For each climate variable, predictors included current-year T_1–3_ and T_4–6_, P_1–3_ and P_4–6_, and D_1–3_ and D_4–6_, together with previous-year T_7–9_ and T_10–12_, P_7–9_ and P_10–12_, and D_7–9_ and D_10–12_. Together, these windows span July of the previous year through June of the growth year, a period over which climate–tree-growth relationships are often strongest (44, 30).

Ring-width chronologies carry not only a climate signal but also biological persistence, disturbance, and other non-climatic variation (12). The context predictors **z***_c_* (species, elevation, and chronology identity) absorb persistent non-climatic differences between chronologies, so that the learned climate response is not confounded by them, and the lag-1 term *y_c,t_*_−1_ captures the strong year-to-year autocorrelation of ring width (44). Chronology and species identity were encoded as categorical predictors using XGBoost’s native categorical handling rather than as ordinal numeric variables.

The data were divided chronologically into training (1902–1980), validation (1982–1990) and testing (1992–2024) sets, with lagged predictors constructed within each split. Species were retained only when represented by at least 10 training chronologies, 5 post-1980 chronologies, 200 training observations and 100 post-1980 observations. Model fitting was restricted to Northern Hemisphere chronologies, where the network is concentrated, so that the calendar-based predictor windows followed a common annual cycle. Applying the same criteria to the Southern Hemisphere retained only 69 chronologies and 840 observations after 1990, with records ending in 2012, so the Southern Hemisphere was excluded. We evaluated 48 randomly sampled hyperparameter combinations, using early stopping within each trial and selecting the model by validation RMSE. The selected model used 1,431 boosting rounds, with all fitted hyperparameters reported in Supplementary Table S2.

### Model validation and characterization

Held-out predictive skill was evaluated on the 1992–2024 test set, where the model achieved *R*^2^ = 0.257, Pearson *r* = 0.511 and RMSE = 0.257 across 50,849 chronology-year observations from 3,165 chronologies (Fig. S8). Because the model also has access to chronology identity, species, elevation, and previous-year growth, a concern is that its apparent skill could come from this non-climatic structure — learning each chronology’s typical growth level or its year-to-year persistence — rather than from a genuine climate–tree-growth relationship. Since the projections rest entirely on the learned climate response, we tested this directly with a climate-row permutation test. For each permutation, the 12 climate predictors were shuffled together as an intact block across training rows, while chronology identity, species identity, elevation and lagged growth were left unchanged. Shuffling the predictors as a block, rather than independently, preserves the realistic covariance among temperature, precipitation and VPD across their windows, so each climate vector remains physically plausible; only its association with the corresponding chronology-year growth observation is destroyed. We refitted the model for each of 1,000 permutations using the same selected hyperparameters and 1,431 boosting rounds. The resulting null distribution had mean test *R*^2^ = 0.023 (s.d. 0.003); none of the 1,000 permutations reached the observed *R*^2^ = 0.257, giving empirical *P <* 0.001 (Fig. S9). The model’s skill therefore depends on a genuine learned climate–tree-growth relationship, which the non-climatic features cannot reproduce on their own.

On the test set, precipitation is the model’s largest single contributor by mean absolute SHAP (SHapley additive explanations) value (32), more than twice either temperature or VPD, followed by lag-1 growth, VPD, temperature, and the context predictors (Fig. S10). Our analysis, however, is based on the baseline-relative change in attribution (ΔSHAP; below) rather than absolute importance, and the two need not agree: precipitation dominates the model’s present-day predictions, whereas the projected late-century attribution change is led by previous-year maximum temperature (T_7–9_) and current-year VPD (D_4–6_) (Fig. S7).

### Projecting climate-driven growth responses

Projecting ring-width chronologies forward poses two structural difficulties. First, chronologies are stationary by construction: de-trending removes low-frequency growth trends, so a chronology carries no projectable trend and its expected level is fixed rather than drifting. Second, ring-width series are strongly autoregressive, and the model uses previous-year growth *y_c,t_*_−1_ as a predictor; under future climate no observed lag-1 value exists, and feeding the model its own predictions would either compound error or, given stationarity, collapse the series toward its mean. Naive year-by-year forecasting of RWI is therefore ill-posed.

We instead isolate the climate-driven component of the response by neutralizing the autoregressive predictor. For chronology *c*, year *t*, scenario *s* and climate model *m*, the neutral-history prediction was

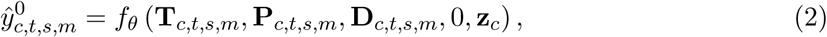

where the lag-1 term is set to 0, corresponding to RWI = 1, the stationary expectation of a detrended chronology. This uses only the climate predictors and the known chronology context **z***_c_* to evaluate the learned climate response for a given chronology, with the chronology, species, and elevation terms held at their trained values. Neutralizing the lag-1 term does not distort this response: in the held-out SHAP audit, previous-year growth was nearly uncorrelated with the model’s climate predictors (maximum |*r*| = 0.052), and a Savitzky–Golay smoothing of its SHAP dependence crossed zero at RWI*_t_*_−1_ = 0.969, close to the neutral value RWI*_t_*_−1_ = 1, so the term contributes negligibly there (Fig. S11). Because the neutral-history prediction does not depend on the chronology having been observed in a given year, it is applied consistently across the full 1902–2100 period for every chronology in the network.

All reported values are changes relative to each chronology’s own historical baseline rather than absolute predicted RWI. For a target window *W*, we compared the neutral-history prediction over that window with the corresponding prediction over the baseline period *B* (1902–1980):

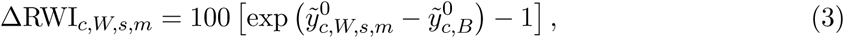

where *ỹ*^0^ denotes the period median of annual neutral-history predictions of *ŷ*^0^. The exponential converts log-RWI differences to proportional changes, so all results are reported as percentage change in RWI relative to baseline (% ΔRWI). The baseline period coincides with the model training period, where the representation of each chronology’s climate response is most strongly constrained; projections measure departure from this grounded reference, including where climate moves away from the training distribution. Most results are reported at the grid-cell level. Chronologies were assigned to 1^◦^ × 1^◦^ latitude–longitude grid cells; within each cell and climate model we took the median chronology change over the target window, and for projected periods then took the median across models, giving one value per occupied cell. Reporting one value per occupied cell keeps the maps and distributions spatially balanced rather than dominated by densely sampled regions (Fig. S12). Projected windows were 30 years long, advancing from the observed period into the CMIP6 projections; the eight-decade baseline and the multi-year windows together suppress interannual noise. Because the autoregressive channel is neutralized and chronology context is held fixed in both terms, the baseline and target predictions differ only in their climate inputs, so ΔRWI isolates the effect of climate change on the modeled growth response.

### Within-chronology validation

Beyond overall predictive skill, we asked a more specific question: within a single chronology, does the model correctly tell good growth years from bad ones on the basis of climate? This is a stronger test than fitting across sites, because it asks whether the model has learned each chronology’s own year-to-year response to climate. For each test-period chronology, we used the neutral-history predictions to identify its years of high and low climate-driven growth, taking the upper and lower quartiles as the model’s favorable and unfavorable years. If the model has captured a real climate response, the tree should have grown more in those favorable years than in the unfavorable ones; we therefore compared the observed growth difference between the two sets of years with the predicted difference, for chronologies with at least ten test-period years. Across 2,541 chronologies, the observed and predicted contrasts were correlated (Pearson *r* = 0.80) and agreed in sign for 86% of chronologies, far above the 50% expected by chance (1,000-permutation null; *P <* 0.001). The observed contrasts were on average 1.5 times larger than predicted (fitted slope 1.54; Fig. S13), so the model captures the pattern of climate-driven growth while understating its amplitude. Defining favorable and unfavorable years at other percentile thresholds gave similar results.

### Out-of-domain climate screening

Projected climates can move into combinations of conditions not represented in the training data, where the model would extrapolate (52). To identify these, we screened each prediction against the training climate cloud in the joint space of the twelve climate predictors (three variables, each summarized over four three-month periods; chronology context and lag-1 growth were not screened) (37). We standardized the twelve predictors over the training data and, in this standardized space, computed for each prediction its Euclidean distance to its *k*-th nearest training neighbor (*k* = 20). A prediction was flagged as out-of-domain (OOD) when this distance exceeded the 99th percentile of the corresponding neighbor distances among the training data itself. Because the screen operates on the joint climate vector rather than on individual variables, it flags novel *combinations* of otherwise-sampled conditions, not only values beyond each variable’s range.

We aggregated this row-level screen to grid cells in the same rolling 30-year windows used for the response projections. For each scenario, climate model, window, and 1^◦^ × 1^◦^ cell, we computed the fraction of prediction rows that were OOD and took the median of this fraction across CMIP6 models; a cell was classified as out-of-domain when this model-median exceeded 50%, so that a cell was excluded only when most models agreed its projected climate lay outside the training climate domain. Observed CRU climate was screened through 2024 and the bias-corrected CMIP6 climate from 2025 onward, so windows spanning the transition combined both; the out-of-domain fraction grew through the century, most steeply under SSP5-8.5 (Fig. S4).

Predictions in out-of-domain cells were withheld from the reported responses, so that projected changes are shown only where the model interpolates within its training climate.

### Statistical support

We estimated the uncertainty of each cell’s ΔRWI with a temporal bootstrap (16). We drew 5,000 bootstrap replicates. In each replicate, we resampled the years of the baseline window and of the 30-year target window, with replacement, and applied the same resampled years to every chronology. From these resampled years we recomputed, in order: each chronology’s baseline and target-window medians, each chronology’s ΔRWI, the median across chronologies within each 1^◦^ × 1^◦^ grid cell, and the median across the twelve climate models. The bootstrap therefore captures the temporal sampling uncertainty of a finite window, while the chronologies and models contributing to each cell stay fixed. For windows spanning the observed-to-projected transition, we resampled the CRU years once per replicate and shared them across models, and resampled the CMIP6 years separately within each model.

From each cell’s 5,000 resampled ΔRWI values, we computed a two-sided bootstrap *P* -value for a change away from zero. We controlled the false discovery rate (FDR) across cells with the Benjamini–Hochberg procedure (5), applied separately within each scenario and window and over in-domain cells only, having excluded out-of-domain cells beforehand. We treated cells with *q* ≤ 0.05 as significant, and took the direction of each significant change from the sign of its point estimate. Out-of-domain cells were held at their last in-domain state for the rolling transition and regime analyses. This single cell-level procedure underlies the results in Figures 2, 3 and 4.

### Within-model spatial spread

The projected response distribution widens spatially through the century (Fig. 2). To confirm this broadening reflects a signal shared across climate models rather than disagreement among them, we recomputed the spatial spread within each model separately. For each CMIP6 model and rolling window, we formed the grid-cell ΔRWI values as above, i.e., the median across chronologies within each in-domain 1^◦^ × 1^◦^ cell, without aggregating across models, and took their interquartile range across space as a measure of spatial spread. We summarized the twelve single-model spatial IQRs by their median and 25th–75th percentile range, and compared them with the spatial IQR of the ensemble-median field used in the main distributions (Fig. S2). This diagnostic uses the same out-of-domain mask as the response projections but no FDR filtering, since it characterizes the full in-domain distribution rather than statistically significant change. The spatial spread increases through the century within individual models, so the broadening does not arise from combining divergent models.

### Regional summaries

Occupied 1^◦^ × 1^◦^ grid cells were assigned to IPCC AR6 WGI reference regions (v4) by their center point, using the land and land–ocean polygons (24). We report the seven represented regions with at least 95 occupied cells, which together hold 956 of the 1,171 occupied cells and 3,650 of the 4,110 chronologies. The threshold falls at a natural break: the smallest retained region has 95 cells and the next below it has 45. Sampling is concentrated in North America and Europe, and no Asian region reached the threshold (Supplementary Table S3).

### Transition and retention analysis

We tracked each cell’s state (positive, negative, or neutral) across the rolling 30-year windows, following how projected growth changes emerged and whether they were retained through to the end of the century. Because change is measured relative to the baseline, cells begin neutral and may depart as the windows advance; for each cell, we recorded the window at which it first became positive or negative and the state of its last in-domain window. Retention was the fraction of cells that first entered a positive (or negative) state and were still in that state at the last window, with cells that never left neutral excluded. We report it as a proportion with a Wilson 95% confidence interval (54) for the full network, and as a rate for each of the seven well-sampled AR6 regions. Cells that left the training climate domain before 2100 were held at their last in-domain state, so that retention was measured over every cell that entered a state, not only those whose climate stayed within the sampled range. Recomputing the analysis with these cells excluded left retention nearly unchanged (negative 89.8% versus 90.1% under SSP2-4.5, 96.7% versus 96.9% under SSP5-8.5).

We examined the regions to test whether the difference between negative and positive retention was systematic rather than the average of a few strongly one-sided regions. A two-sided sign test (binomial, *n* = 7) assessed whether negative retention differed from positive retention across the seven regions; it was higher in all seven (*P* = 0.016). We then estimated the network-wide gap with a two-level bootstrap of 20,000 replicates: each replicate resampled the seven regions and then the cells within each, and we recomputed the entry-weighted negative-minus-positive gap. The empirical 2.5th and 97.5th percentiles of the bootstrap distribution were used as the 95% confidence interval, reflecting both among-region variation and within-region cell-level variation.

Endpoint retention compares cells that entered their state at different times against a common date (2100), so cells entering earlier have longer to depart. To remove this, we also measured retention at matched horizons after entry. For each cell we took its first in-domain, statistically significant non-neutral state and, at each horizon of *h* years, asked whether the cell was still on that same side in the first in-domain state at or beyond *h* years after entry. This measure is observable-only: a cell contributes to horizon *h* only if it has an in-domain state there, and cells drop out as they leave the training climate domain or reach the end of the century, rather than being frozen. Because the observable denominator shrinks with horizon, we show the retention curves only while at least 150 cells of each entry sign remain observable in both scenarios, giving a common horizon of 53 years (Fig. 3c). Negative retention rose and positive retention fell with horizon under both scenarios, widening the gap.

Because this dynamic denominator changes with horizon, we repeated the analysis on a fixed cohort: cells whose first entry could be followed in-domain for the full 50 years after entry, tracked over the same 50-year span. Holding the cohort constant removes any effect of a changing denominator, at the cost of excluding cells that entered too late for a 50-year follow-up before 2100. Within this fixed cohort the retention gap still widened steadily with horizon — from 3 pp after one year to 34 pp by 50 years under SSP2-4.5, and from 5 pp to 50 pp under SSP5-8.5 (Fig. S3) — so the divergence is not an artifact of the shrinking denominator.

### Climate-driver decomposition

To attribute each projected growth change to the underlying climate variables, we decomposed the model’s prediction with SHAP, which distributes a prediction among its input features additively, so that, together with the expected-value term, their contributions sum to the prediction. We write the SHAP value of feature *j* as *ϕ_j_* and computed these with XGBoost’s native tree-based method (33), which is exact and additive on the log(RWI) scale, on the same neutral-history prediction rows used for the projections (lag-1 growth set to zero).

The quantity we attribute is not the prediction itself but its change from baseline. Just as growth change is the predicted RWI in a window relative to the chronology’s baseline, each driver’s contribution is the change in its SHAP value relative to baseline (its ΔSHAP): if a chronology’s predicted growth falls below baseline, we ask how much of that decline each driver accounts for. The decomposition is therefore of the baseline-relative change in attribution, not of absolute SHAP or feature importance.

We grouped the twelve climate features into a *water-supply* contribution, from the four precipitation predictors, and a *warming–demand* contribution, from the four maximum-temperature and four VPD predictors; maximum temperature and VPD were combined because they are strongly correlated and their separate contributions cannot be resolved reliably. For a feature group *G*, the annual SHAP value is the sum over its features, *ϕ_G,c,t,m_* = *_j_*_∈_*_G_ ϕ_j,c,t,m_*, where *c* indexes chronologies, *t* years and *m* climate models (omitted for observed years). As for the growth projections, each group’s ΔSHAP is its value in a *W* = 30-year window relative to the chronology’s *B* = 1902–1980 baseline,

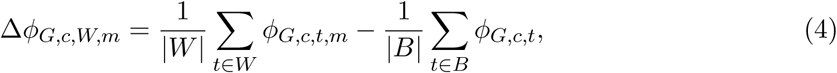

taken as the median across the *M* = 12 climate models, Δ*ϕ_G,c,W_* = median*_m_* Δ*ϕ_G,c,W,m_*. The water–supply, warming–demand and net ΔSHAP were formed on the log scale,

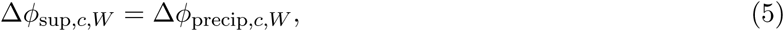

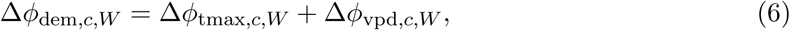

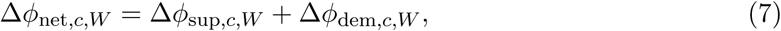

and each converted to a percent-equivalent RWI contribution, matching the units of the growth projections,

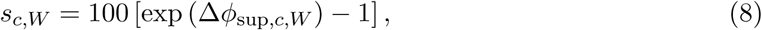

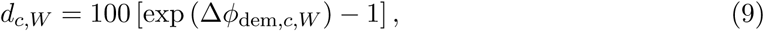

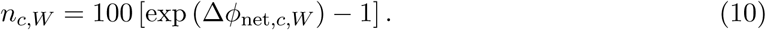

We summarized *s*, *d* and *n* at the 1^◦^ × 1^◦^ cell level as the median across chronologies.

The net contribution *n* is the climate component of the predicted change. It is aggregated on the log-RWI SHAP scale before conversion to percent-equivalent RWI; because the conversion and median aggregation are nonlinear, *n* is not the arithmetic sum of the displayed supply and warming–demand percentages. The static context features are held fixed between baseline and future, but their SHAP values can shift modestly through interactions with the changing climate inputs, so the climate net accounts for most, though not all, of the baseline-relative change. We therefore take each cell’s positive or negative state from the statistically significant change in predicted RWI, and use the decomposition to diagnose the climatic pathway behind it; the sign of the climate net agreed with this state for over 96% of significant cells under both scenarios.

The signs of *s* and *d* place each in-domain, statistically significant cell in the supply–demand plane: co-enhancement (*s >* 0*, d >* 0) and co-suppression (*s <* 0*, d <* 0) where the two contributions share a sign, and supply-driven (*s >* 0*, d <* 0*, n >* 0) or demand-driven (*s >* 0*, d <* 0*, n <* 0) where they oppose, split by the sign of the climate net *n* rather than by the cell’s significant positive or negative state. In-domain cells whose change was not statistically significant form a fifth, neutral regime. The reverse case, a negative supply against a positive demand (*s <* 0*, d >* 0), was negligible here, since the warming–demand contribution is negative across most of the network; the few such cells were neutral. The overall direction of change is statistically significant, but the division within it turns on the smaller contribution, whose sign is not separately tested; we therefore interpret regime composition at the population and regional level rather than as a fixed label for any individual cell.

### Response-surface and attribution analysis

To identify which predictors carried the projected changes, we ranked the twelve climate predictors by the magnitude of their late-century ΔSHAP, summed in absolute value across chronologies (using the median across models, as above). Under SSP2-4.5, four predictors — P_4–6_, D_4–6_, T_7–9_ and T_10–12_ — account for 53% of this summed absolute change in attribution, and we used them for the response-surface analysis (Fig. 5; ranking in Fig. S7). The ranking is similar under SSP5-8.5, with T_10–12_ falling just outside the leading four.

Each response surface shows how predicted growth changes as we vary just two predictors at a time, leaving all the other inputs to the model as they are. The other inputs are supplied by *N_c_* = 350 chronology-year observations drawn at random from the 1902–1980 reference period, and the surface is the prediction averaged over them. We sweep the two focal predictors across a grid of values; at each grid point, we set the two focal predictors to that grid point in every one of these observations and average the model’s predictions:

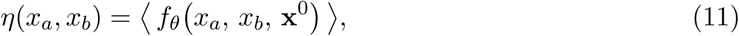

where *x_a_* and *x_b_* are the two focal predictors, **x**^0^ holds the remaining fixed inputs from one reference-period observation — chronology context **z**, neutral lag-1 growth (*y_t_*_−1_ = 0) and the non-focal climate predictors — and ⟨·⟩ averages over the *N_c_* observations. The surface is thus a function of the two focal predictors alone. For visualization, each surface was shifted by subtracting the prediction at the middle grid cell of that two-predictor panel; thus the color scale shows relative changes in predicted RWI across the plotted climate plane, not absolute predicted RWI or change relative to the historical baseline.

This surface is a scalar field *η* over climate space, and projected climate change acts on it as a displacement field Δ**q**: each chronology is carried from its baseline position (start) to its future position (end). The impact of climate change is the change in the scalar field along this displacement,

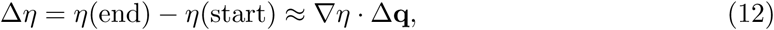

so it depends on both the local gradient of the surface and the direction and length of the displacement. This makes impact distinct from sensitivity, the gradient alone: a location may sit on a steep part of the surface yet move little, or move far along a nearly flat direction. We evaluated the displacement field by binning the projected movements of the underlying chronologies in baseline climate space and, on each surface, show the mean displacement per bin together with the response change it produces (Fig. 5). The displacement field is a summary: the same point in climate space can be occupied by chronologies from different regions projected to experience different climate change, so their displacements differ, and the underlying unit remains the individual chronology.

Because the surfaces are evaluated over a full grid, they extend into climate combinations the model was not trained on, where predictions flatten as the model saturates rather than extrapolating steeply (consistent with the out-of-domain screening above). We therefore overlay the observed baseline climate states on each surface, so the sampled region of climate space — where the surface is supported by data — is visible, and the surface should not be read where no points fall.

The main text shows selected surfaces; the full set of pairwise surfaces among the twelve predictors, and regional breakdowns of the leading pairs, are shown in Figs. S14–S24.

## Data and code availability

The tree-ring chronologies analysed in this study are publicly available from the International Tree-Ring Data Bank (39) and the European beech network of Klesse et al. (29). Observed and projected climate data are publicly available from CRU TS v4.09 (21) and NEX-GDDP-CMIP6 (48, 47), respectively. The derived data underlying each figure, together with the code to reproduce the analyses and figures, will be made available on GitHub and archived on Zenodo with a permanent DOI upon publication.

## Supplementary Information

### S1 Supplementary Equations

#### vapor pressure deficit calculation

Vapor pressure deficit (VPD) was calculated from maximum temperature and atmospheric vapor pressure. Saturation vapor pressure was computed using a Bolton-style formulation,

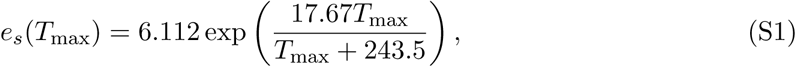

where *T*_max_ is monthly maximum temperature in ^◦^C and *e_s_* is the saturated atmospheric vapor pressure in hPa. For observed CRU TS data, VPD was then calculated as

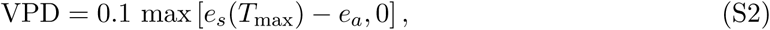

where *e_a_* is CRU TS vapor pressure in hPa and the factor 0.1 converts hPa to kPa.

For NEX-GDDP-CMIP6 projections, daily VPD was calculated from daily maximum temperature and near-surface relative humidity RH as

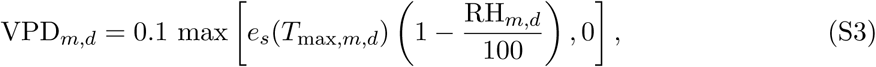

where *m* indexes climate models and *d* indexes days. Relative humidity was clipped to the interval 0–100% before calculating VPD. Daily VPD values were averaged to monthly means before bias correction and seasonal aggregation.

### S2 Supplementary Tables and Figures

**Table S1:** CMIP6 models used in the projection ensemble. The twelve CMIP6 general circulation models whose NEX-GDDP-CMIP6 downscaled projections were used under SSP2-4.5 and SSP5-8.5. Variant labels give the CMIP6 realization-initialization-physics-forcing member used. Projected growth responses are summarized as the median across these models.

| Model | Variant |
| --- | --- |
| ACCESS-CM2 | r1i1p1f1 |
| CanESM5 | r1i1p1f1 |
| CNRM-CM6-1 | r1i1p1f2 |
| EC-Earth3 | r1i1p1f1 |
| FGOALS-g3 | r3i1p1f1 |
| GFDL-ESM4 | r1i1p1f1 |
| IPSL-CM6A-LR | r1i1p1f1 |
| MIROC6 | r1i1p1f1 |
| MPI-ESM1-2-LR | r1i1p1f1 |
| MRI-ESM2-0 | r1i1p1f1 |
| NorESM2-MM | r1i1p1f1 |
| UKESM1-0-LL | r1i1p1f2 |

**Table S2:** Model inputs, data partitioning and fitted XGBoost configuration. Predictors, chronological train/validation/test split, and the hyperparameters of the selected XGBoost model, chosen by validation RMSE from 48 sampled configurations (Methods). Predictor windows are three-month summaries denoted by their first month.

| Item | Detail |
| --- | --- |
| Target | Annual ring-width index (RWI), log-transformed. |
| Climate predictors | **Maximum temperature:** current-year $T_{1-3}$ and $T_{4-6}$; previous-year $T_{7-9}$ and $T_{10-12}$.<br>**Precipitation:** current-year $P_{1-3}$ and $P_{4-6}$; previous-year $P_{7-9}$ and $P_{10-12}$.<br>**VPD:** current-year $D_{1-3}$ and $D_{4-6}$; previous-year $D_{7-9}$ and $D_{10-12}$. |
| Context predictors | Chronology identity, species identity, elevation and previous-year log RWI. |
| Data partitioning | **Training:** 1902–1980; 278,691 observations (77.2%).<br>**Validation:** 1982–1990; 31,385 observations (8.7%).<br>**Testing:** 1992–2024; 50,849 observations (14.1%). |
| Boosting | 1,431 rounds; learning rate = 0.030608. |
| Tree structure | Histogram tree method; maximum depth = 8; minimum child weight = 8; maximum bin count = 512. |
| Sampling | Row subsample = 0.878678; column subsample = 0.737010. |
| Regularization | $L_1 = 1.223213$; $L_2 = 0.769702$; $\gamma = 0.015007$. |
| Objective and seed | Squared-error objective; seed = 20260442. |

**Table S3:** AR6 regional magnitude summary for the late-century period (2071–2100). For each represented IPCC AR6 reference region: the number of occupied 1^◦^ grid cells and the number of chronologies they contain, and, for each scenario, the percentage of mapped cells outside the training climate domain (OOD), the median predicted change in ring-width index relative to the 1902–1980 baseline (%ΔRWI), and the number of in-domain statistically significant cells with negative and positive change (neg./pos.). A dash indicates no in-domain cells remained. The seven regions with at least 95 occupied cells (bold) are summarized in the main text.

| Region | Name | Cells | Chronologies | SSP2-4.5 |  |  | SSP5-8.5 |  |  |
| --- | --- | --- | --- | --- | --- | --- | --- | --- | --- |
|  |  |  |  | OOD% | Median | neg./pos. | OOD% | Median | neg./pos. |
| **WNA** | W. North America | **185** | 917 | 15.1 | −0.4 | 27/17 | 45.9 | −0.9 | 17/11 |
| **WCE** | W. & C. Europe | **161** | 1042 | 1.9 | −2.5 | 87/4 | 7.5 | −6.2 | 127/2 |
| **NWN** | N.W. North America | **151** | 417 | 6.6 | −0.8 | 59/20 | 12.6 | −3.1 | 92/15 |
| **ENA** | E. North America | **130** | 374 | 10.0 | −0.1 | 13/28 | 29.2 | −0.9 | 33/15 |
| **MED** | Mediterranean | **126** | 414 | 4.0 | −4.2 | 107/1 | 51.6 | −7.7 | 61/0 |
| **NEU** | N. Europe | **108** | 224 | 3.7 | −0.0 | 31/29 | 4.6 | −1.6 | 52/23 |
| **CNA** | C. North America | **95** | 262 | 25.3 | −1.2 | 12/0 | 65.3 | −3.8 | 20/0 |
| NCA | N. Central America | 45 | 129 | 68.9 | −10.7 | 11/0 | 95.6 | −8.5 | 2/0 |
| ESB | E. Siberia | 37 | 44 | 0.0 | −4.6 | 26/0 | 10.8 | −7.4 | 28/0 |
| NEN | N.E. North America | 25 | 48 | 0.0 | −3.6 | 23/0 | 0.0 | −5.4 | 24/0 |
| TIB | Tibetan Plateau | 22 | 62 | 27.3 | 0.3 | 3/6 | 50.0 | 2.7 | 0/8 |
| EEU | E. Europe | 21 | 34 | 9.5 | −6.1 | 16/0 | 52.4 | −8.2 | 8/0 |
| RAR | Russian Arctic | 16 | 22 | 0.0 | 0.2 | 3/3 | 0.0 | −0.8 | 6/2 |
| ECA | E. Central Asia | 13 | 27 | 0.0 | −0.3 | 2/5 | 7.7 | −2.8 | 7/3 |
| WCA | W. Central Asia | 12 | 47 | 0.0 | −0.9 | 5/0 | 25.0 | −4.4 | 6/1 |
| WSB | W. Siberia | 10 | 19 | 10.0 | −1.8 | 5/0 | 10.0 | −3.5 | 9/0 |
| RFE | Russian Far East | 5 | 11 | 0.0 | −1.7 | 4/0 | 20.0 | −3.9 | 4/0 |
| SAS | S. Asia | 4 | 5 | 25.0 | −0.2 | 1/1 | 50.0 | 0.4 | 0/1 |
| SCA | S. Central America | 4 | 11 | 75.0 | −10.2 | 1/0 | 100.0 | — | 0/0 |
| EAS | E. Asia | 1 | 1 | 0.0 | 0.2 | 0/0 | 0.0 | 1.8 | 0/1 |

**Figure S1:**
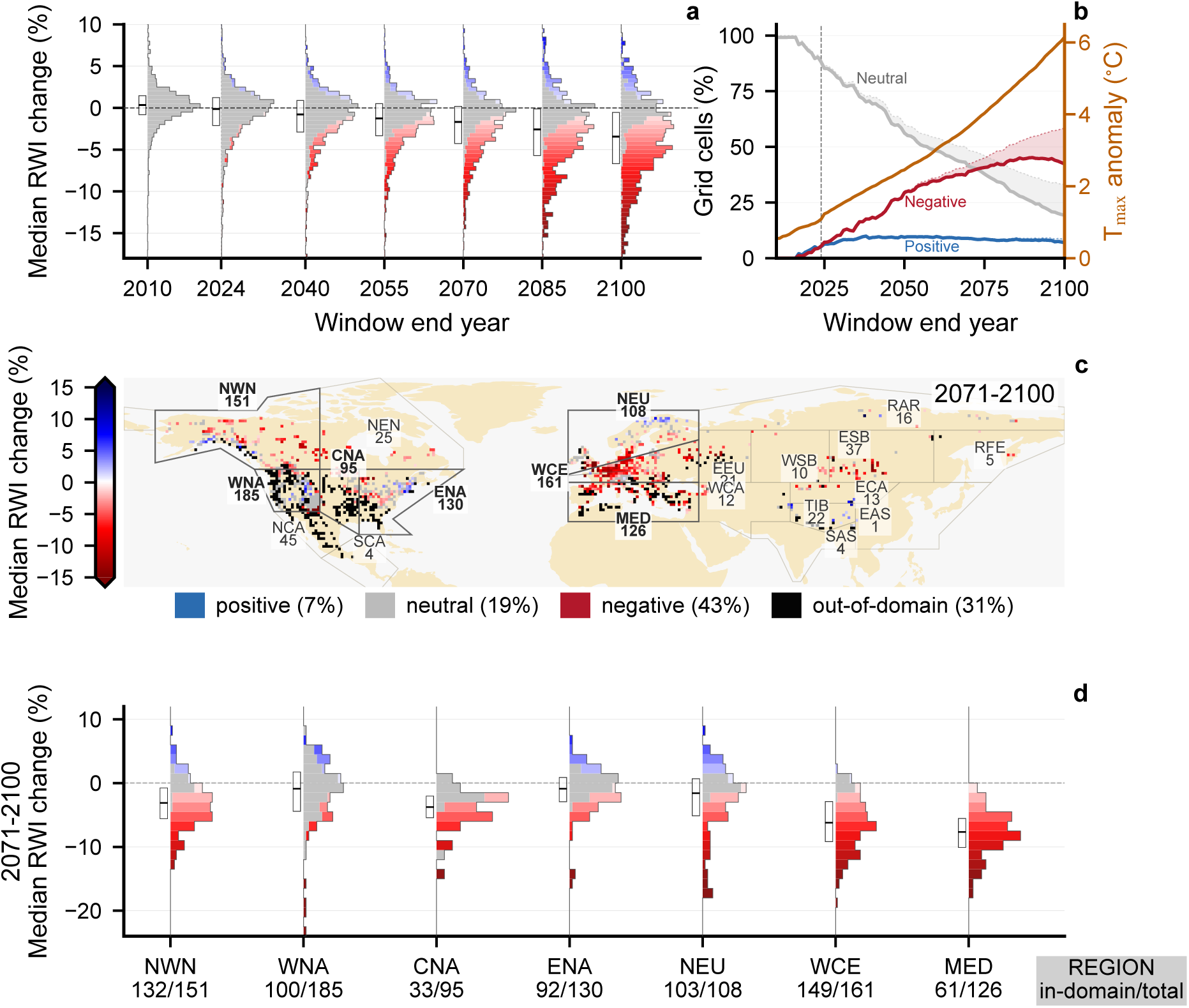
Projected declines are amplified under the high-forcing SSP5-8.5 scenario. As Fig. 2, but for SSP5-8.5. The decreases and broadening are stronger than under SSP2-4.5: the median falls further, the upper quartile drops below zero, and out-of-domain climate becomes more widespread by late century. Panels, encoding, and the temperature scale match Fig. 2.

**Figure S2:**
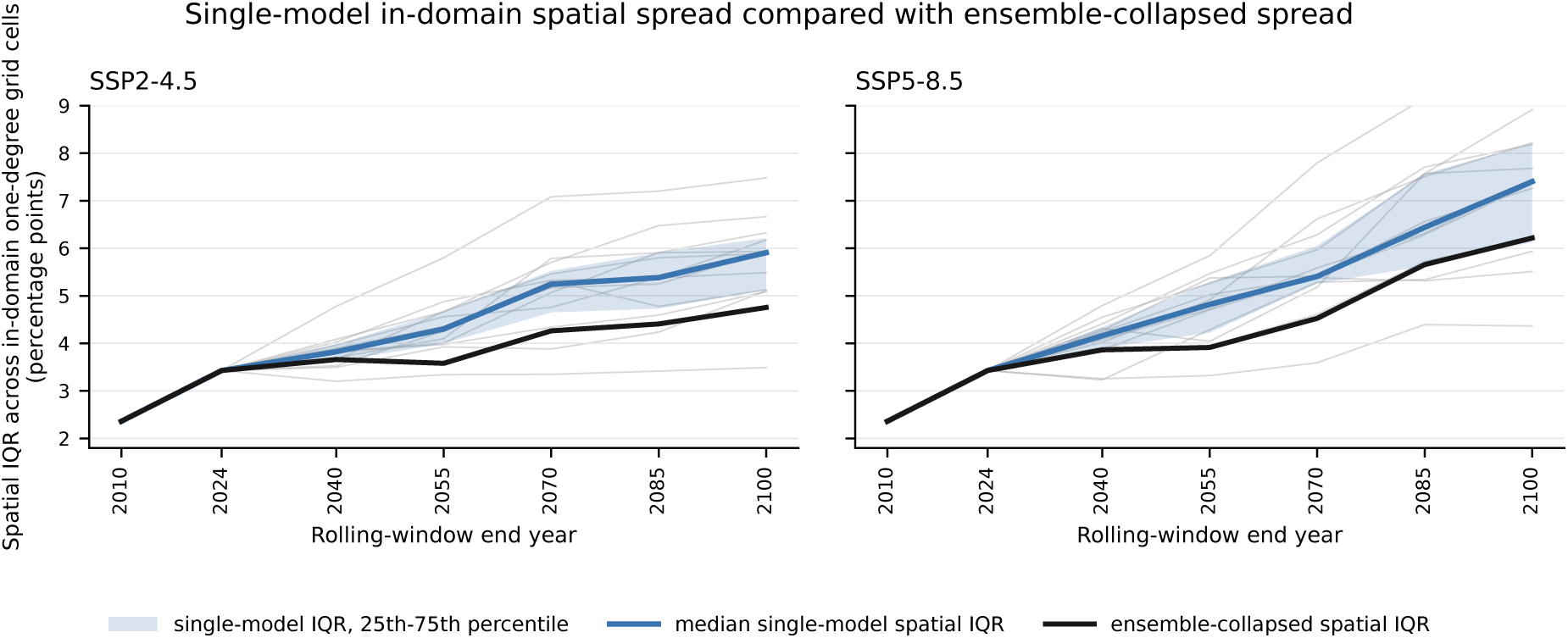
The projected response distribution broadens within individual CMIP6 models. Spatial interquartile range (IQR) of predicted RWI change across in-domain 1^◦^ grid cells, in the same rolling 30-year windows as Fig. 2, for SSP2-4.5 and SSP5-8.5. For each model, grid-cell change is the median predicted RWI change relative to the 1902–1980 baseline after aggregating chronologies within cells; the spatial IQR is then taken across cells. Thin gray lines are the individual CMIP6 models, the blue line their median and the blue band their 25th–75th percentile range, and the black line is the ensemble-median field used in Fig. 2a. Cells outside the climate domain are excluded using the same mask as Fig. 2; no FDR filtering is applied, so the panel describes the full in-domain distribution. The spatial spread increases through the century within individual models under both scenarios, so the broadening in Fig. 2a does not arise from combining divergent models into the ensemble.

**Figure S3:**
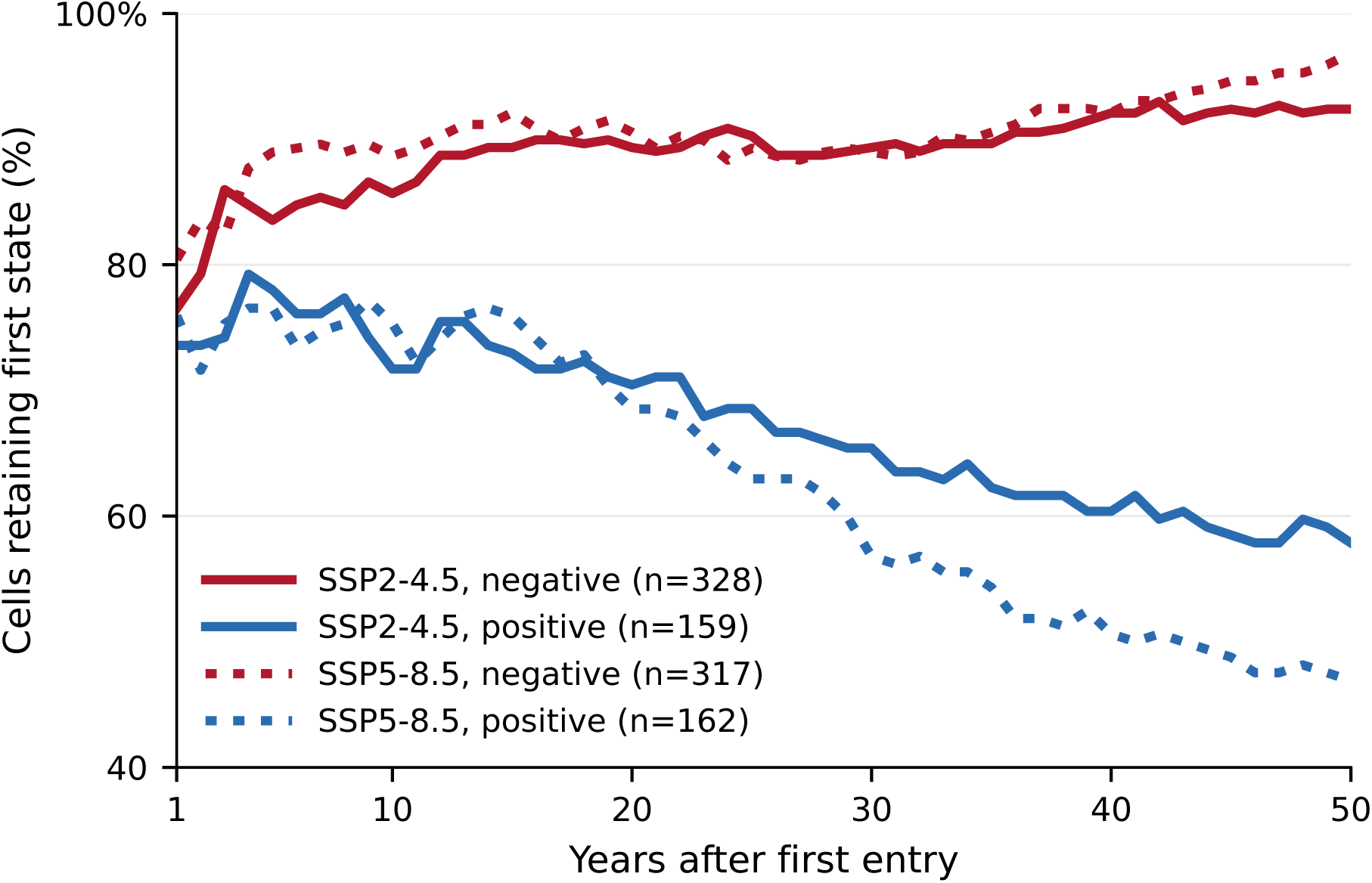
The retention asymmetry holds within a fixed 50-year cohort, ruling out a denominator artifact. Fraction of cells still on their first-entry side as a function of time since first entry, for positive and negative entries, restricted to the fixed cohort of cells that could be followed in-domain for the full 50 years after entry. Unlike the dynamic matched-horizon curves in Fig. 3c, the same cells are used at every horizon from 1 to 50 years, so the denominator does not change; out-of-domain states are not frozen, and cells entering too late for a 50-year follow-up before 2100 are excluded. Line style distinguishes the two scenarios, as indicated in the legend. Within this constant cohort the gap between negative and positive retention still widens with time since entry, reaching 34 and 50 percentage points by 50 years under SSP2-4.5 and SSP5-8.5, reproducing the divergence of the dynamic analysis and confirming it is not driven by the changing denominator.

**Figure S4:**
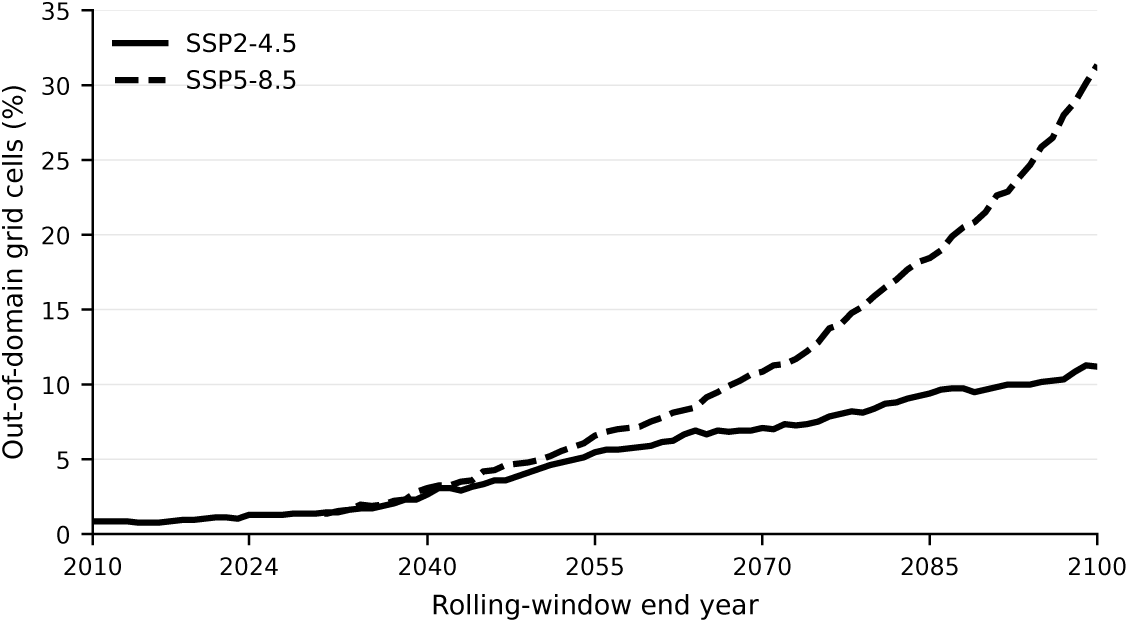
An increasing fraction of the network moves outside the training climate domain, especially under SSP5-8.5. Percentage of mapped 1^◦^ grid cells classified as out-of-domain, in the same rolling 30-year windows as Fig. 2, using the same climate-space mask, which flags cells whose projected climate is poorly represented in the training data. The solid line is SSP2-4.5 and the dashed line SSP5-8.5. The out-of-domain fraction stays low through the observed CRU TS period and rises under the CMIP6 projections, reaching its largest values late in the century under SSP5-8.5.

**Figure S5:**
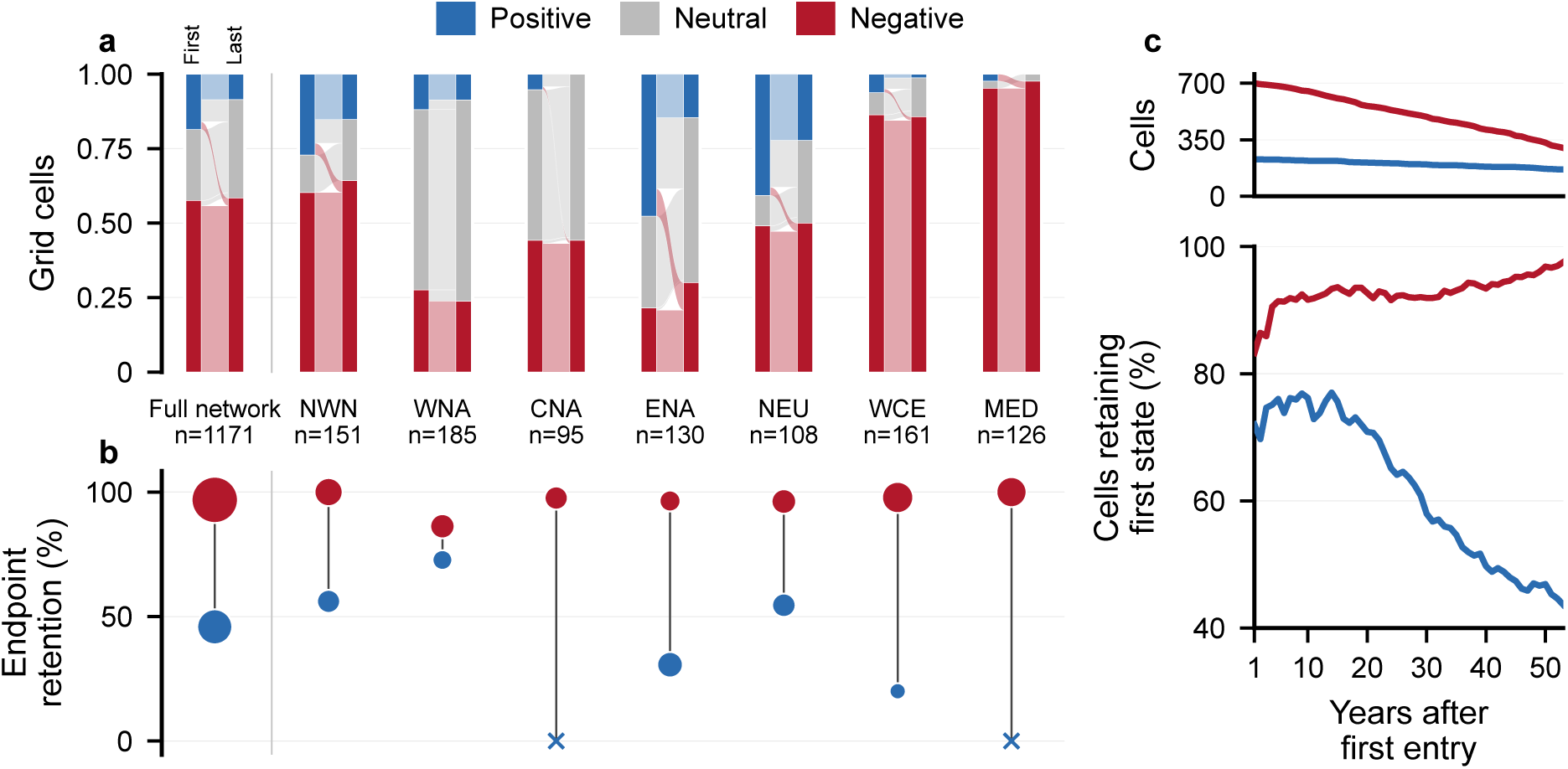
The retention asymmetry strengthens under the high-forcing SSP5-8.5 scenario. As Fig. 3, but for SSP5-8.5. The gap between negative and positive retention widens: almost no negative state reverts, while a larger share of positive states fades to neutral, and downward crossings between signs increase. Panels, state definition, and encoding match Fig. 3.

**Figure S6:**
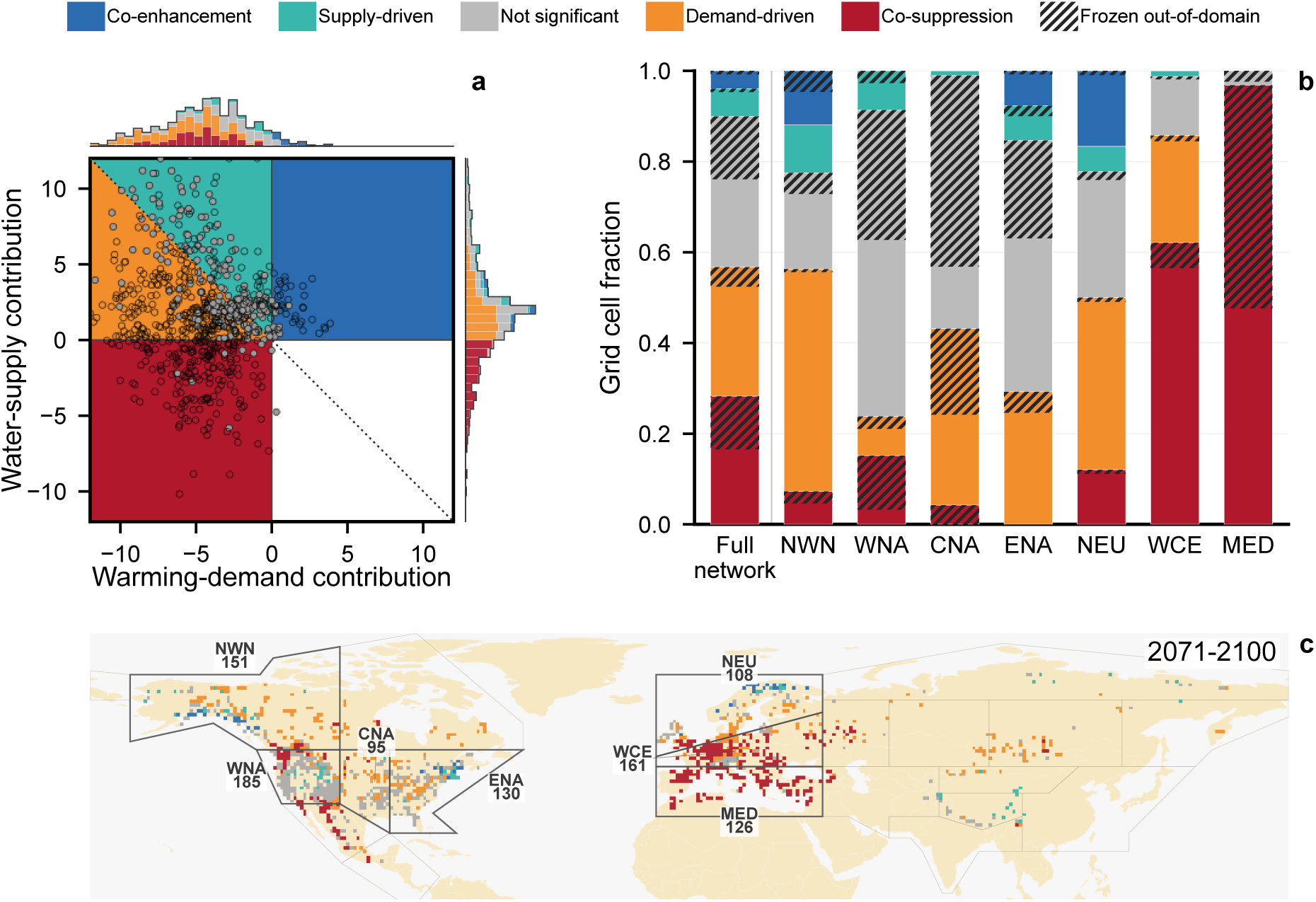
Under SSP5-8.5, surviving gains depend increasingly on water supply as the warming–demand contribution turns more negative. As Fig. 4, but for SSP5-8.5. The warming–demand contribution is negative over a larger share of the network, and among the responses that remain positive, the supply-driven share rises at the expense of co-enhancement, so gains rest increasingly on water supply alone. Panels, driver decomposition, regime definitions, and encoding match Fig. 4.

**Figure S7:**
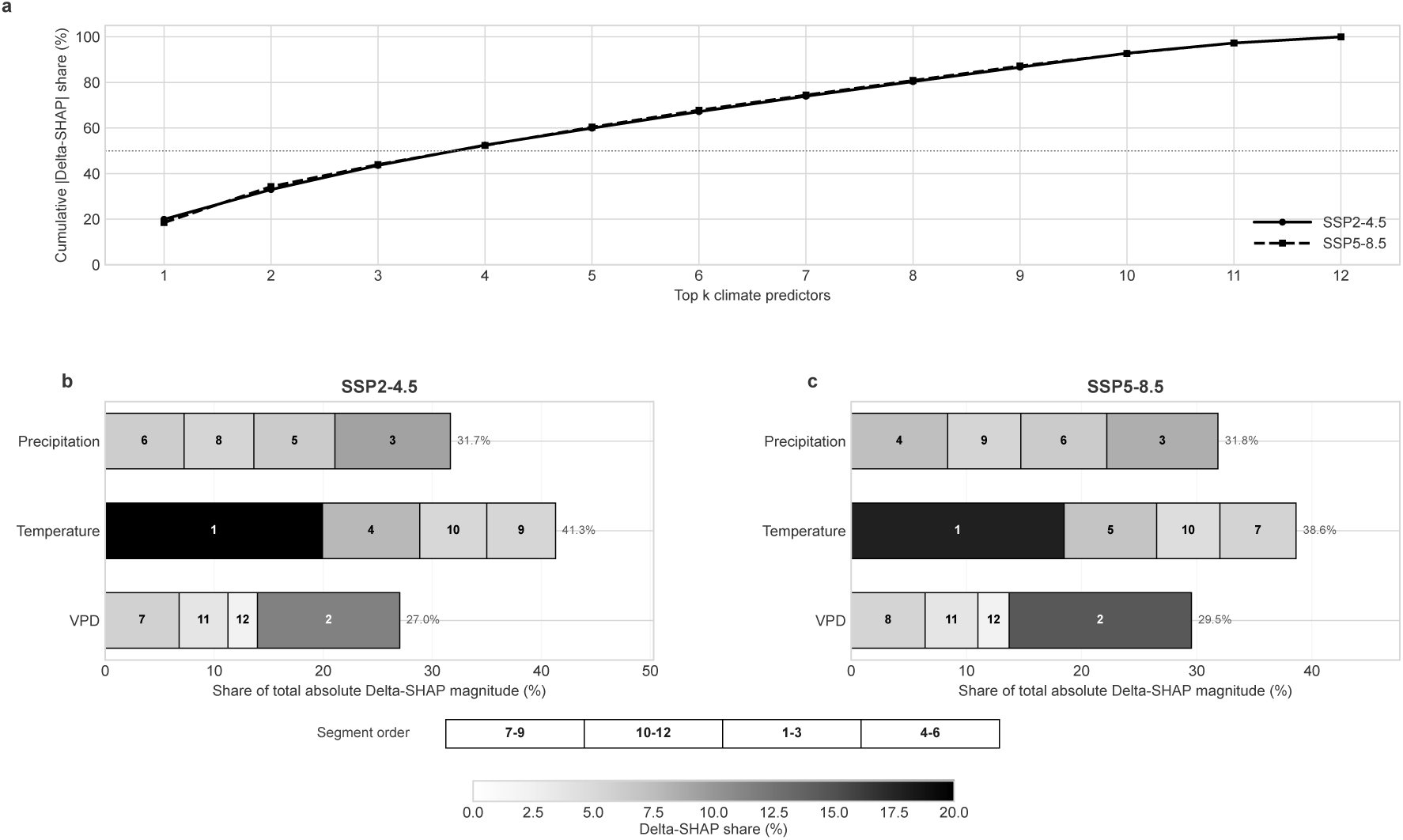
Projected changes in climate attribution are led by maximum temperature and VPD. For each climate predictor, the late-century ΔSHAP is the baseline-relative change in its SHAP contribution, summed in absolute value across chronologies (median across models). Unlike the absolute importance in Fig. S10, which reflects the fitted model as it stands, this measures which predictors drive the *change* in growth under future climate. (a) Cumulative share of the total absolute ΔSHAP captured by the top-ranked predictors, for SSP2-4.5 and SSP5-8.5. The four leading predictors account for about half of the total under both scenarios (53% under SSP2-4.5, 52% under SSP5-8.5), and the top eight for about 80%. (b, c) Family-level composition for SSP2-4.5 and SSP5-8.5. Each bar sums the four windows of a predictor family, with blocks ordered by window as in the legend; numbers inside blocks give each predictor’s rank by total absolute ΔSHAP. Maximum temperature contributes the largest share of the projected change, followed by precipitation and VPD, with previous-year T_7–9_ the single largest predictor under both scenarios.

**Figure S8:**
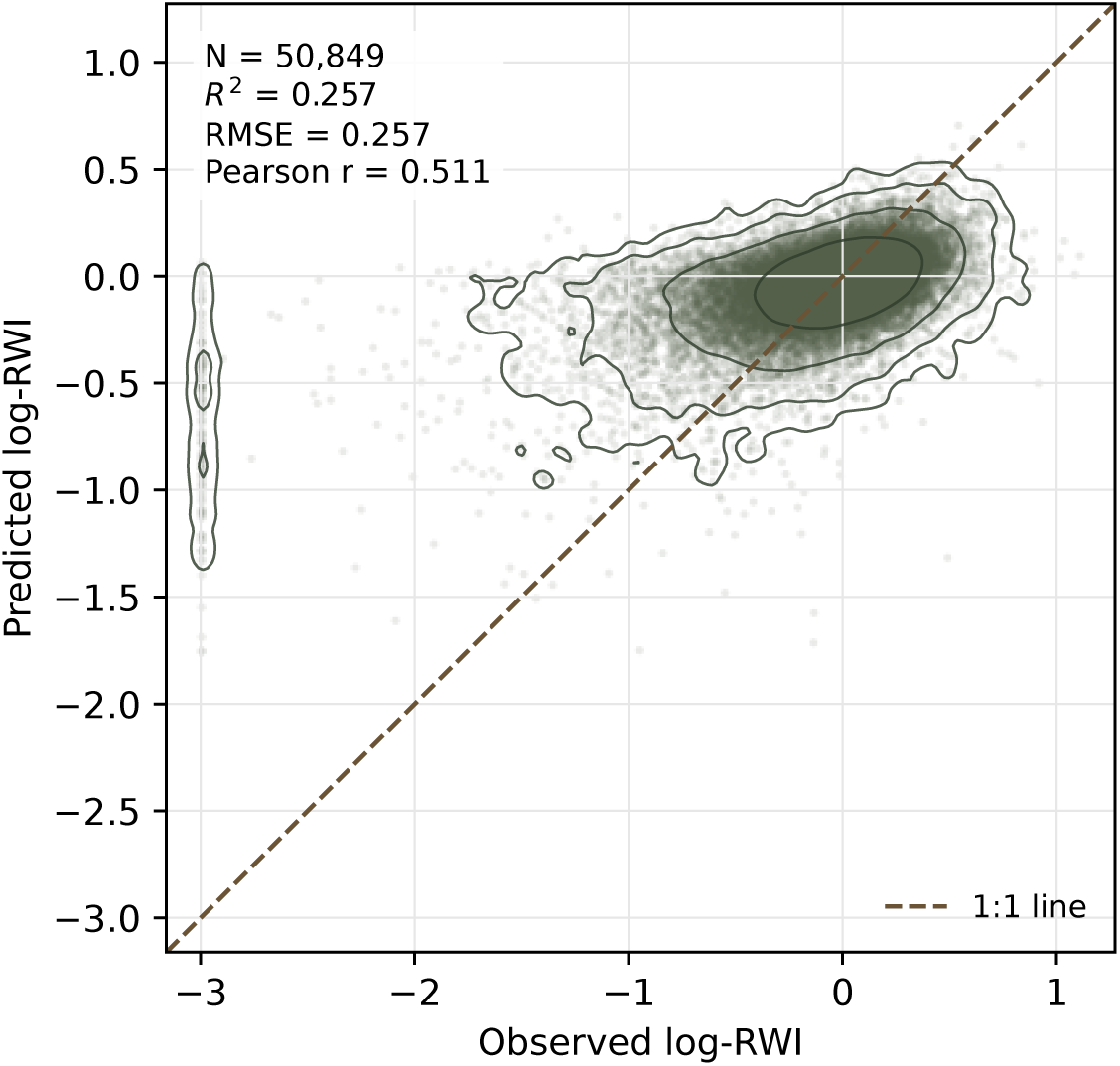
Held-out observed versus predicted growth. Each point is one held-out chronology-year observation from the 1992–2024 test set: the *x* axis is observed log-RWI and the *y* axis is model-predicted log-RWI for the same observation. The dashed line marks one-to-one agreement, and contour lines show the density of observations in the point cloud. The column of observed values near log(RWI) ≈ −3 corresponds to near-zero RWI observations set to the floor RWI = 0.05 before the log transform (Methods). Across 50,849 held-out observations, the model explained 25.7% of annual log-RWI variation (*R*^2^ = 0.257; RMSE = 0.257; Pearson *r* = 0.511).

**Figure S9:**
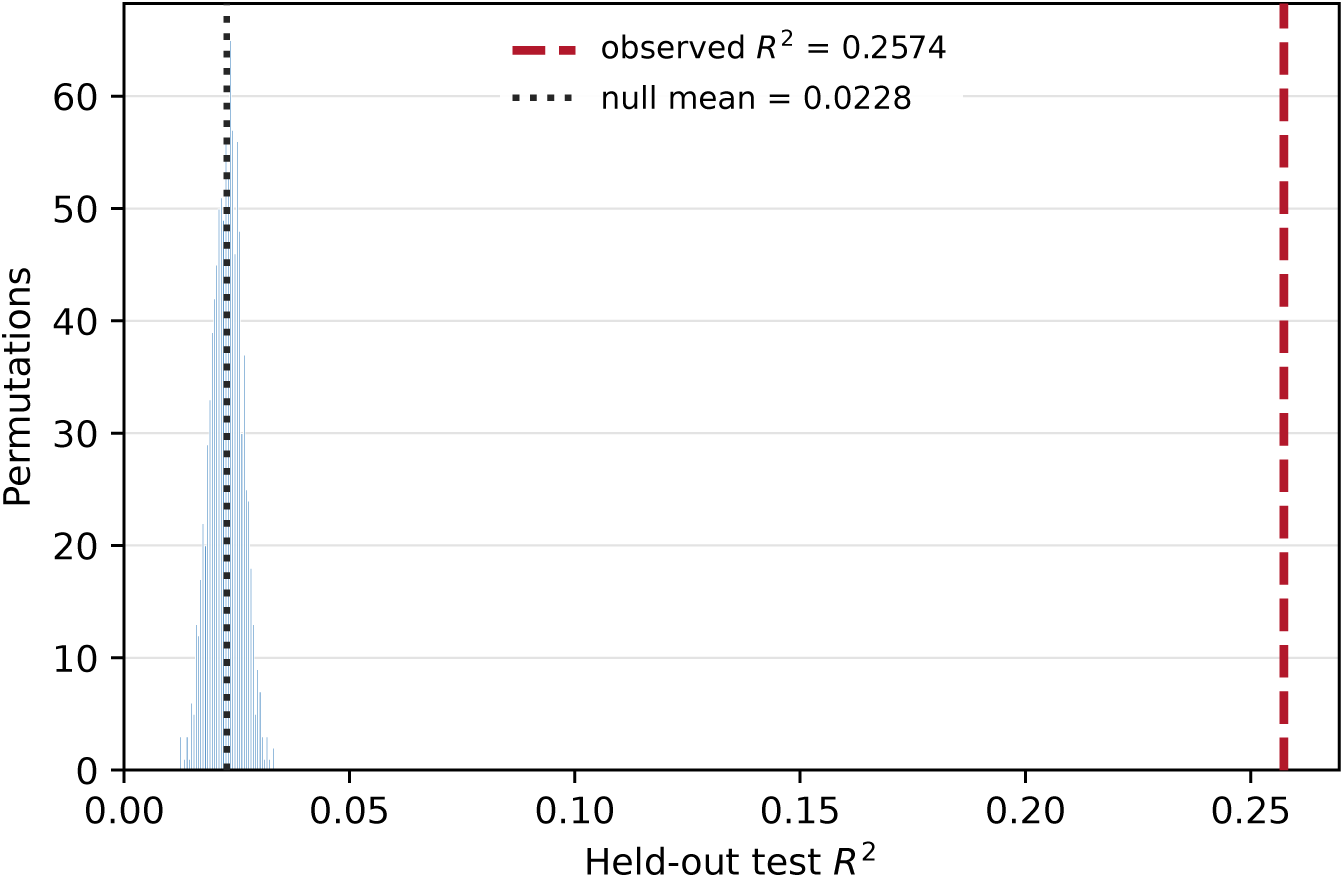
Predictive skill rests on a learned climate response. The 12 climate predictors were permuted together as intact climate rows, preserving realistic within-year covariance among them while removing their correspondence to each chronology-year’s growth; the context predictors (species, elevation and chronology identity) and lag-1 growth were left untouched. For each permutation the full model was retrained on the permuted data, so each of the 1,000 nulls is a model given every opportunity to fit the shuffled dataset. The null distribution had mean test *R*^2^ = 0.023 (s.d. 0.003), whereas the observed model achieved *R*^2^ = 0.257, and no permutation approached the observed value (*P <* 0.001). Skill emerges only when climate and growth retain their real correspondence, showing that the model’s predictions rest on a learned climate response.

**Figure S10:**
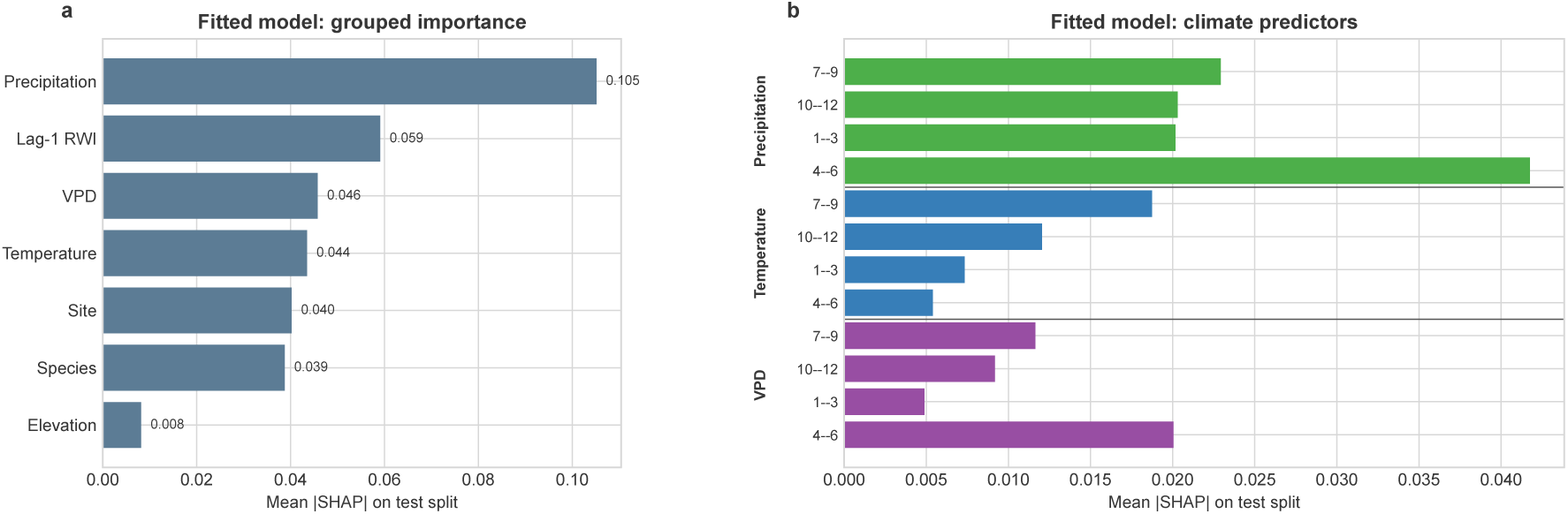
Precipitation is the model’s largest contributor to fitted-model predictions. We measure each predictor’s importance on the held-out test split as its mean absolute SHAP value — how much, on average, it moves the model’s prediction. This importance is a property of the fitted model as it stands, and is distinct from the projected *change* in attribution (ΔSHAP; Fig. S7) analyzed in the main results: one measures which predictors the model relies on, the other which predictors drive the change under future climate. (a) Importance by predictor, with the twelve climate predictors summed within their precipitation, maximum-temperature and VPD families and shown alongside lag-1 growth and the context predictors (chronology identity, species and elevation). Precipitation is the largest single contributor, followed by lag-1 growth. (b) Importance of the twelve individual climate predictors, grouped by family with the four windows of each in calendar order.

**Figure S11:**
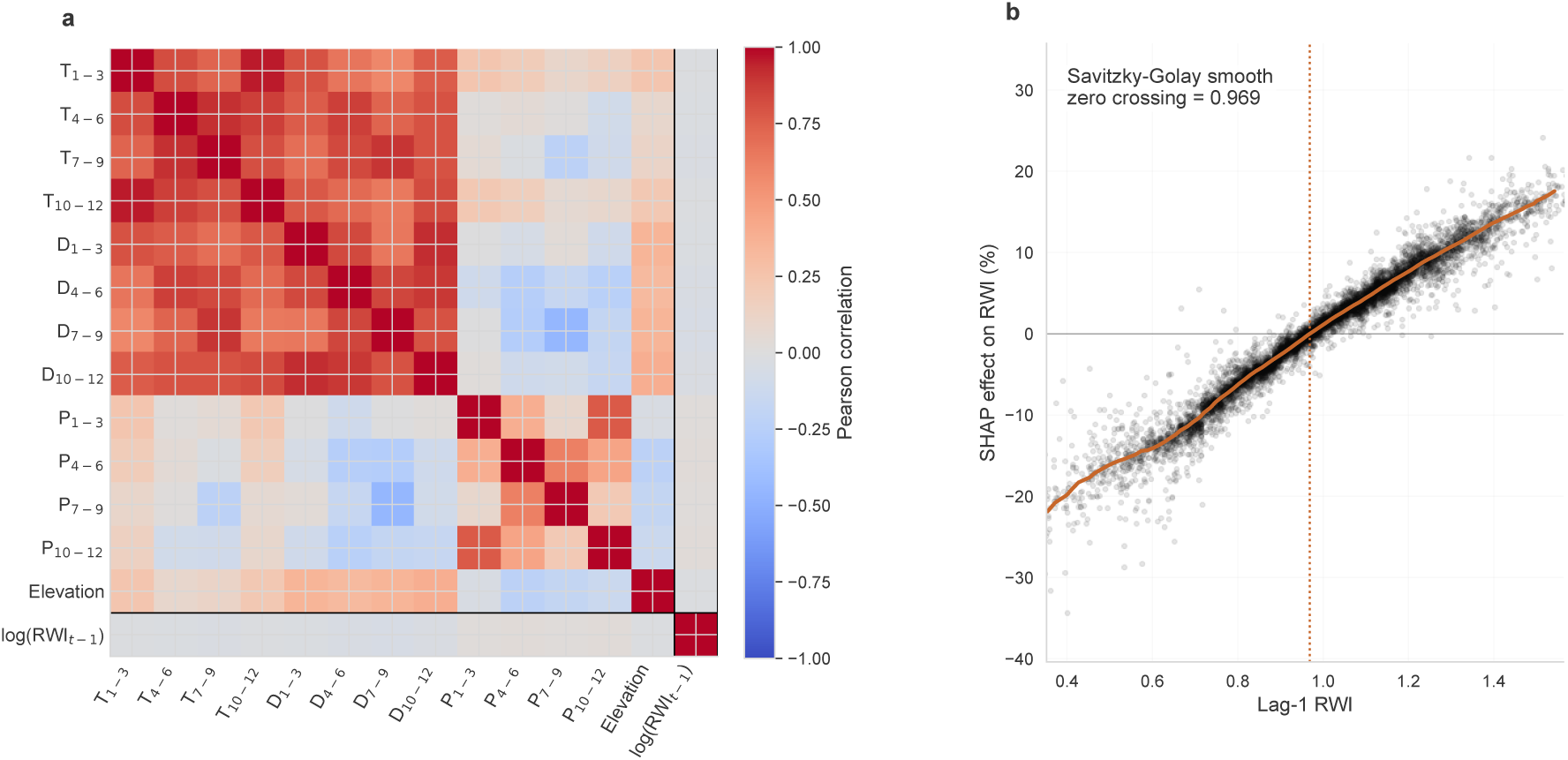
Lag-1 growth is weakly correlated with climate predictors and has near-neutral SHAP contribution at typical previous-year growth. These two properties support setting lag-1 growth to its neutral value when projecting the climate response (Methods, “Projecting climate-driven growth responses”). (a) Ordered Pearson correlation matrix for the numeric model predictors. The outlined row and column mark previous-year log RWI; its correlations with the 12 climate predictors are small, with maximum |*r*| = 0.052. (b) Held-out SHAP dependence for the lag-1 RWI term. Points are individual held-out observations, and the orange curve is a Savitzky–Golay smooth of binned median SHAP effects. The smooth crosses zero at RWI*_t_*_−1_ = 0.969, close to the neutral value RWI*_t_*_−1_ = 1, so the lag-1 term contributes negligibly near neutral previous-year growth.

**Figure S12:**
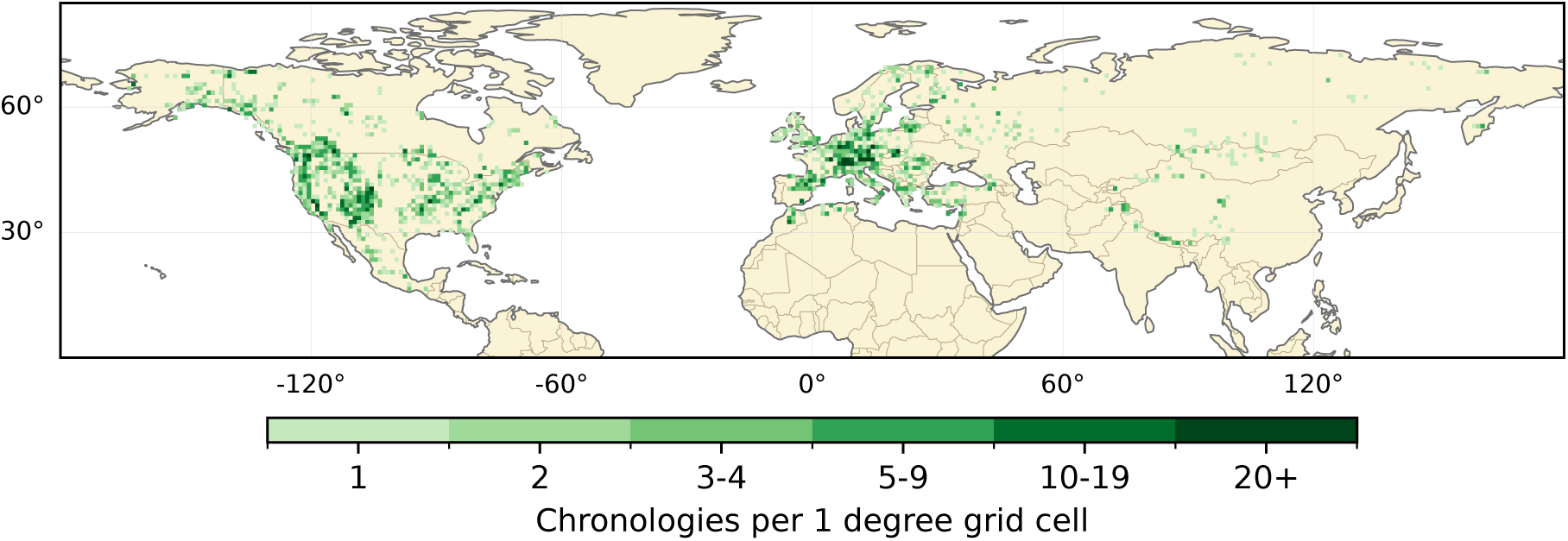
The chronology network is geographically uneven, concentrated in North America and Europe. Number of the 4,110 study chronologies per 1^◦^ grid cell, with color showing binned chronology counts from sparsely sampled cells to dense clusters. This uneven sampling motivates summarizing results at the grid-cell level, which prevents densely sampled regions from dominating the maps and distributions (Methods).

**Figure S13:**
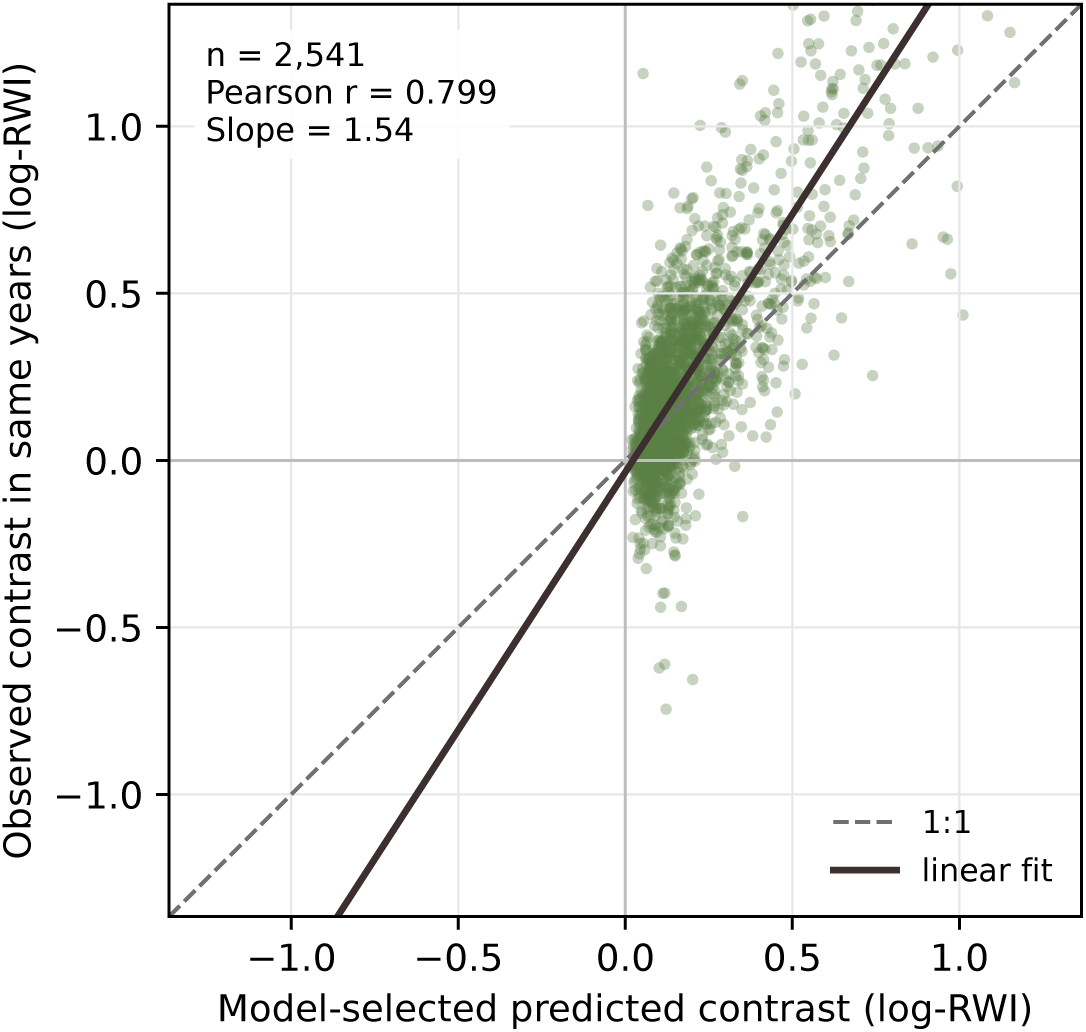
The model reproduces within-chronology growth contrasts but understates their magnitude. This figure tests whether the model can tell a chronology’s good growth years from its bad ones on the basis of climate alone. For each chronology, the model’s climate-driven predictions single out which years should be favorable for growth and which unfavorable. If it has learned a real climate response, the tree should have grown more in the favorable years than in the unfavorable ones. We measure how much growth actually differed between the two sets of years (the observed contrast, *y* axis) and how much the model predicted it should differ (the predicted contrast, *x* axis); each point is one chronology. The dashed line is one-to-one agreement and the solid line the fitted relationship. Across 2,541 chronologies the predicted and observed contrasts are correlated (*r* = 0.80, *P <* 0.001): where the model predicts a larger climate-driven difference between good and bad years, the tree’s growth actually differed more. The fitted slope of 1.54 shows the observed differences exceed those the model predicts, so the model captures the pattern of climate-driven growth but understates its magnitude.

**Figure S14:**
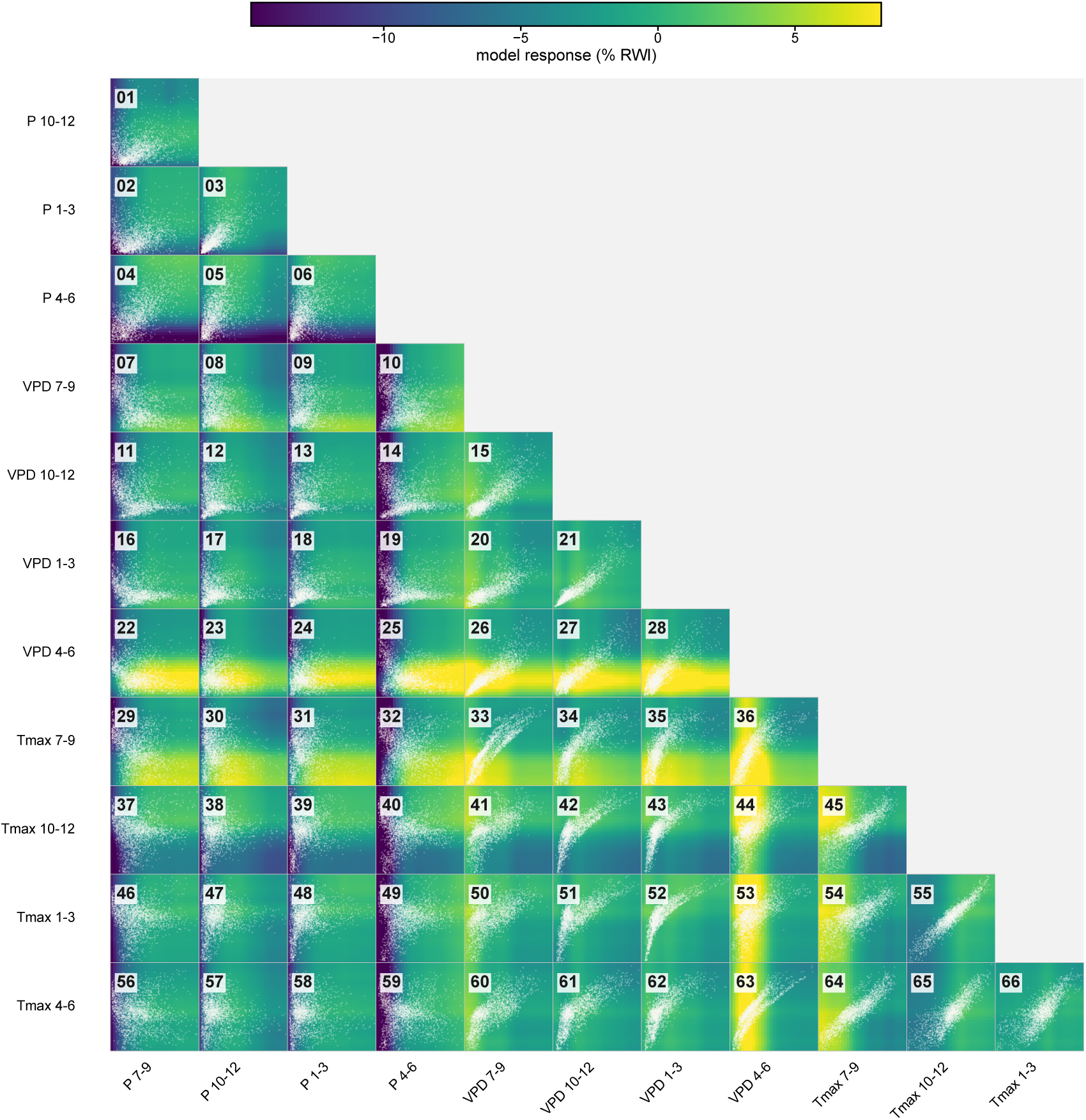
Index of all pairwise climate–tree-growth response surfaces. Thumbnail response surfaces for every pairwise combination of the twelve climate predictors, arranged as a lower-triangle matrix. Each thumbnail shows the model’s learned growth response over that predictor pair (color) with the observed baseline climate states overlaid (points), together indicating which pairwise relationships the model learned and where each is supported by data. The numbered cells index the full-size versions in Figs. S15–S18, where projected climate displacements are added as arrows. Surfaces are read as in Fig. 5.

**Figure S15:**
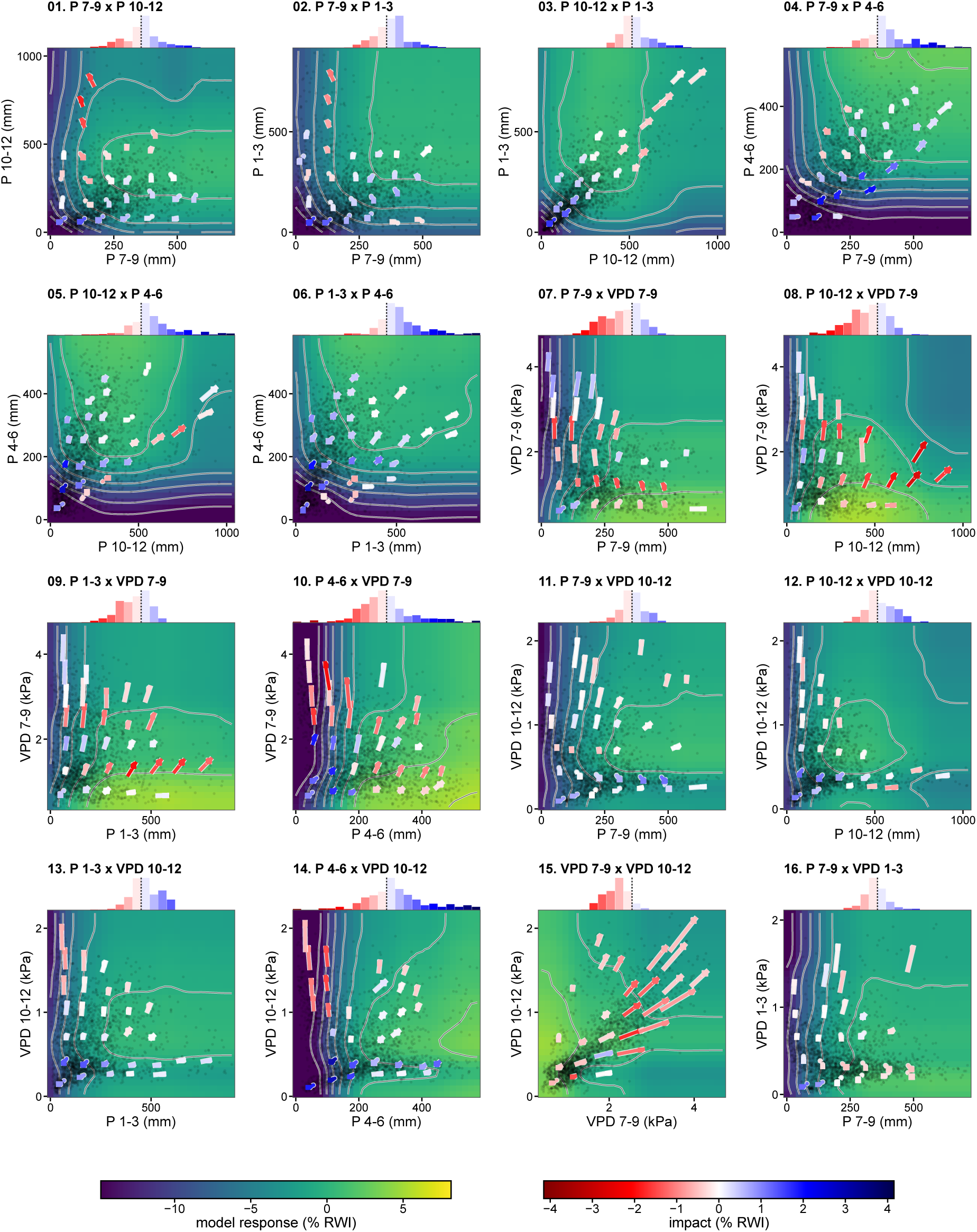
Pairwise climate–tree-growth response surfaces, panels 01–16. Full-size response surfaces for the predictor pairs indexed in Fig. S14. As in Fig. 5, background color is the model’s growth response over the two predictors with contour lines marking levels of equal predicted growth, points are observed baseline climate states marking where the surface is supported by data, and arrows run from the mean baseline to the mean projected late-century climate (SSP2-4.5) of the chronologies in each patch of climate space, colored by the growth change along the move. Histograms above each surface show the distribution of these changes. The surfaces are read through the arrows and how they meet the contours, not the absolute shading.

**Figure S16:**
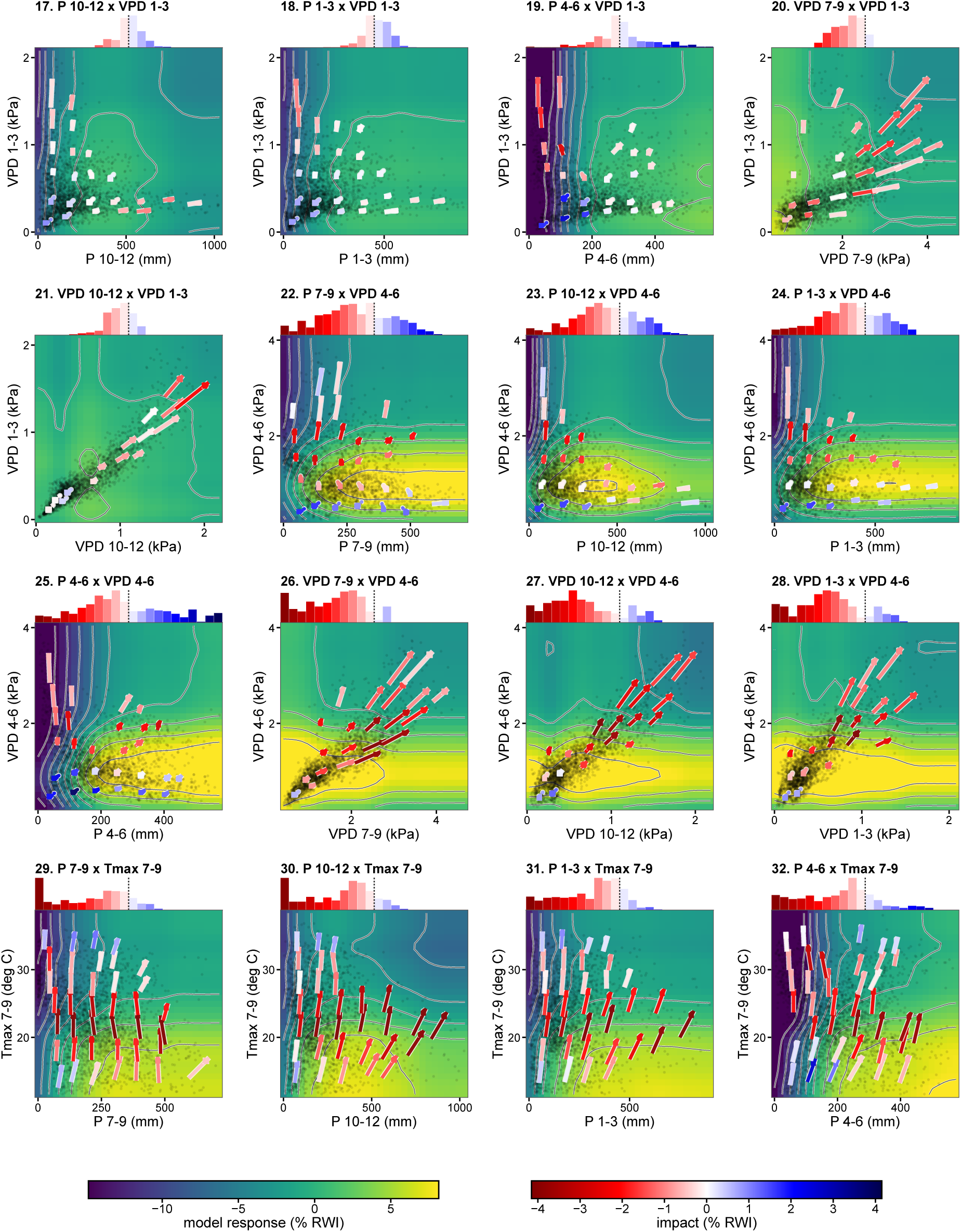
Pairwise climate–tree-growth response surfaces, panels 17–32. Same as Fig. S15, for panels 17–32.

**Figure S17:**
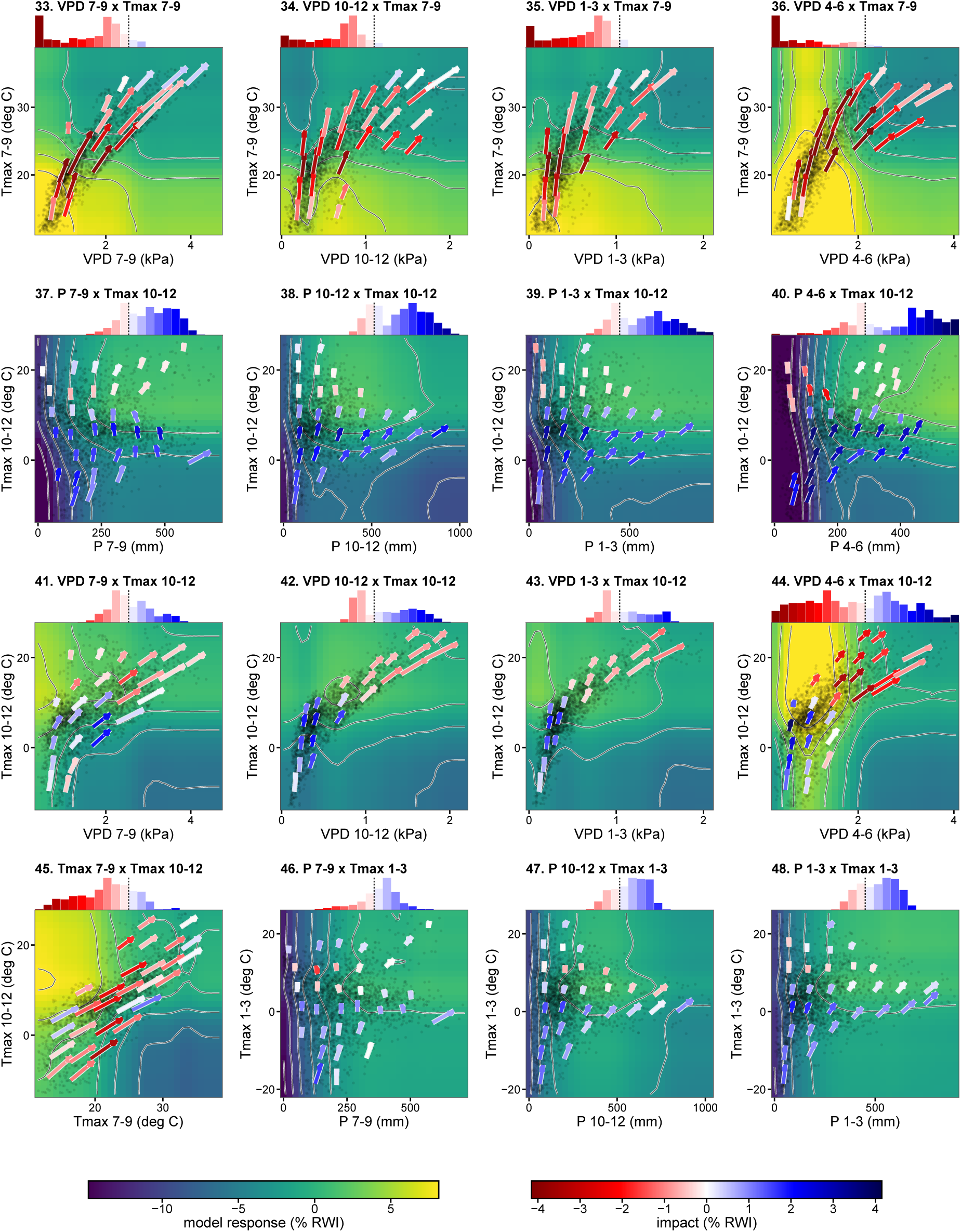
Pairwise climate–tree-growth response surfaces, panels 33–48. Same as Fig. S15, for panels 33–48.

**Figure S18:**
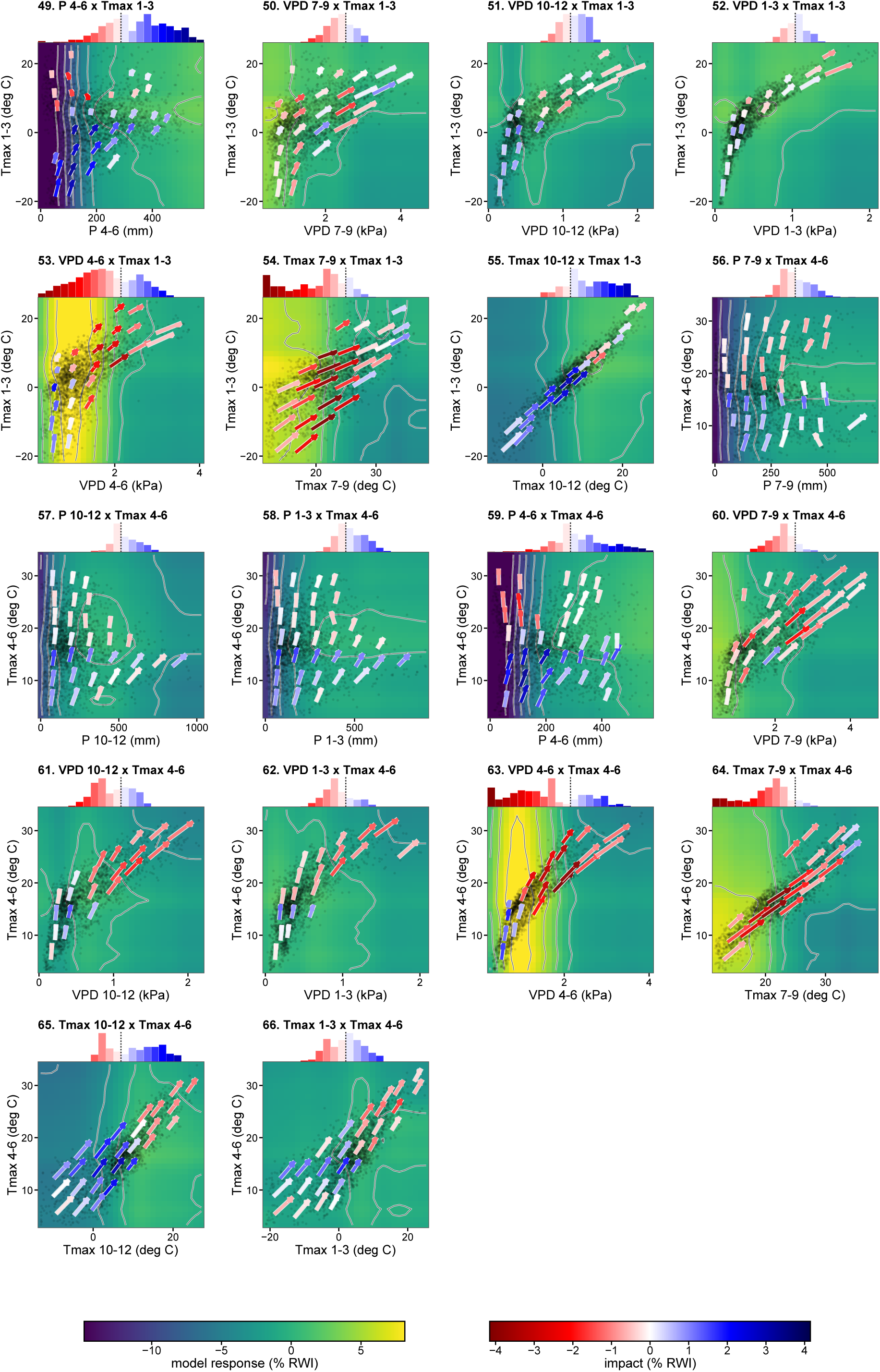
Pairwise climate–tree-growth response surfaces, panels 49–66. Same as Fig. S15, for panels 49–66.

**Figure S19:**
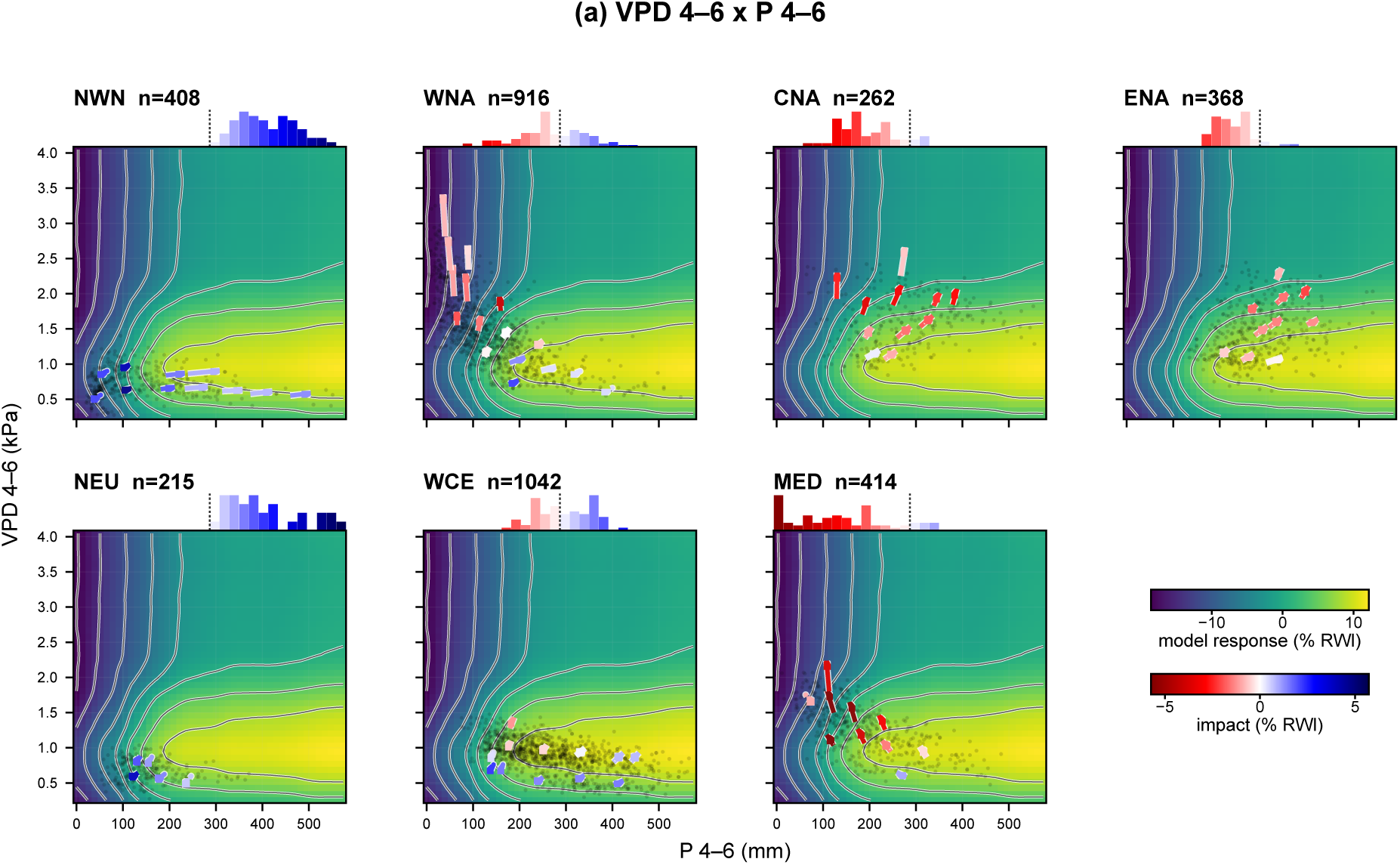
Regional response surfaces for P_4–6_ and D_4–6_. In Fig. 5, only the P_4–6_–D_4–6_ surface (panel a) is broken down by region; the other pairs are shown with their arrows pooled across the network. A pooled arrow summarizes a whole patch of climate space at once, so it can merge chronologies from different regions that are projected to experience different climate change and move in different directions from the same starting point. This set of six figures (Figs. S19–S24) therefore shows all pairwise combinations of the four leading predictors (P_4–6_, D_4–6_, T_7–9_ and T_10–12_) broken down by region, so these region-to-region differences are visible. This figure shows the P_4–6_–D_4–6_ surface for each well-sampled AR6 region, expanding Fig. 5a. Points, arrows and histograms are read as in Fig. S15.

**Figure S20:**
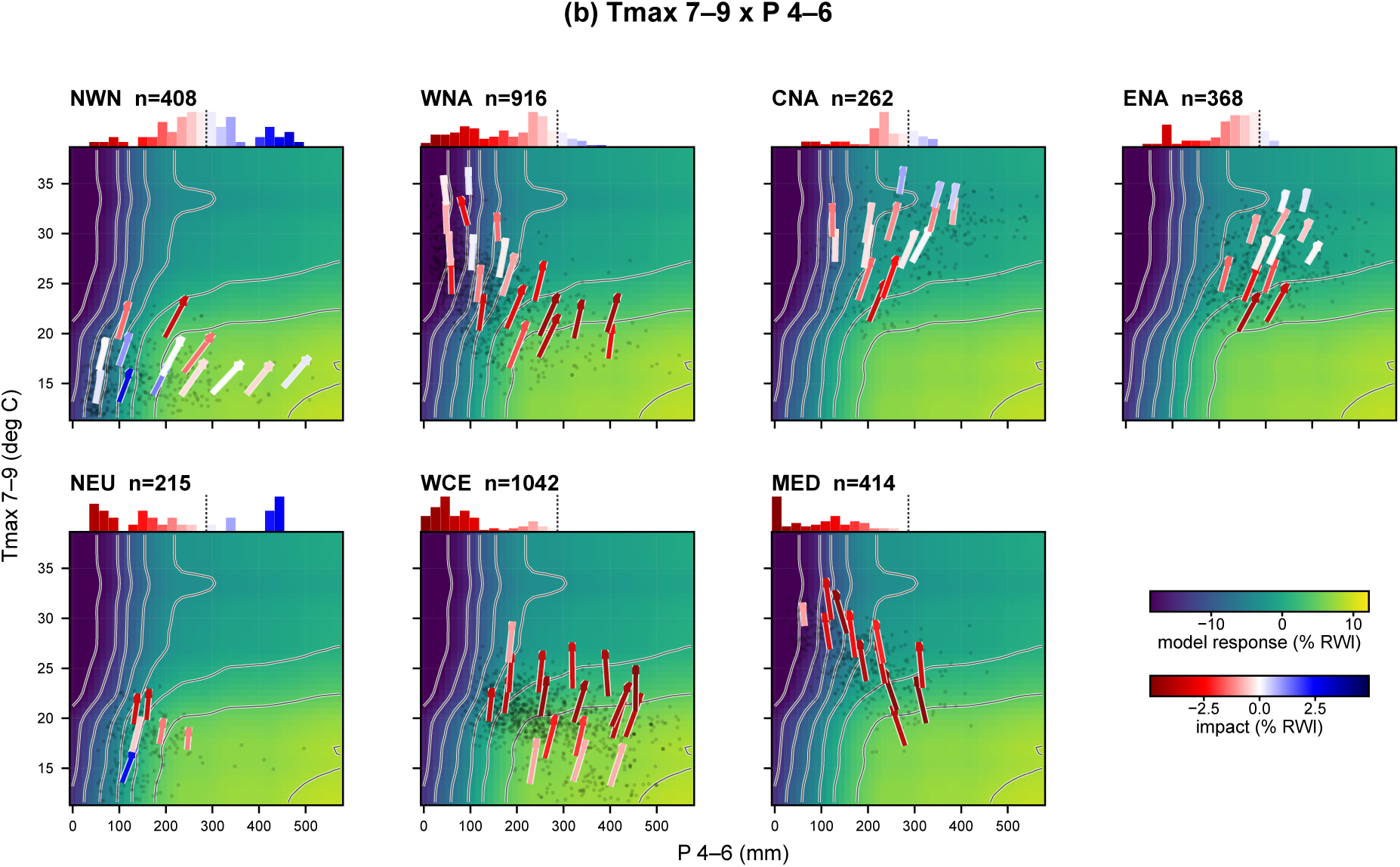
Regional response surfaces for P_4–6_ and T_7–9_. As Fig. S19, for the P_4–6_–T_7–9_ predictor pair.

**Figure S21:**
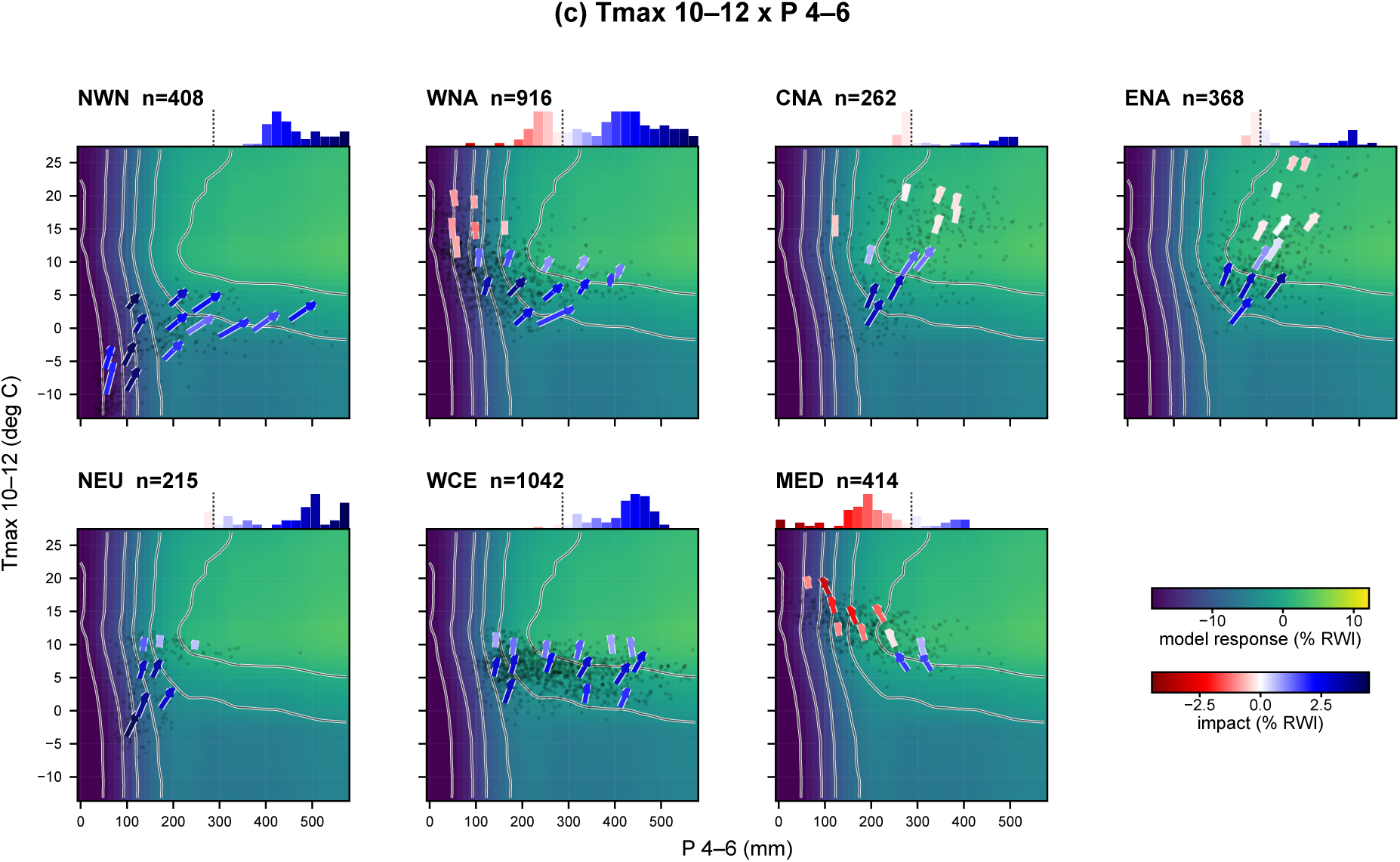
Regional response surfaces for P_4–6_ and T_10–12_. As Fig. S19, for the P_4–6_–T_10–12_ predictor pair.

**Figure S22:**
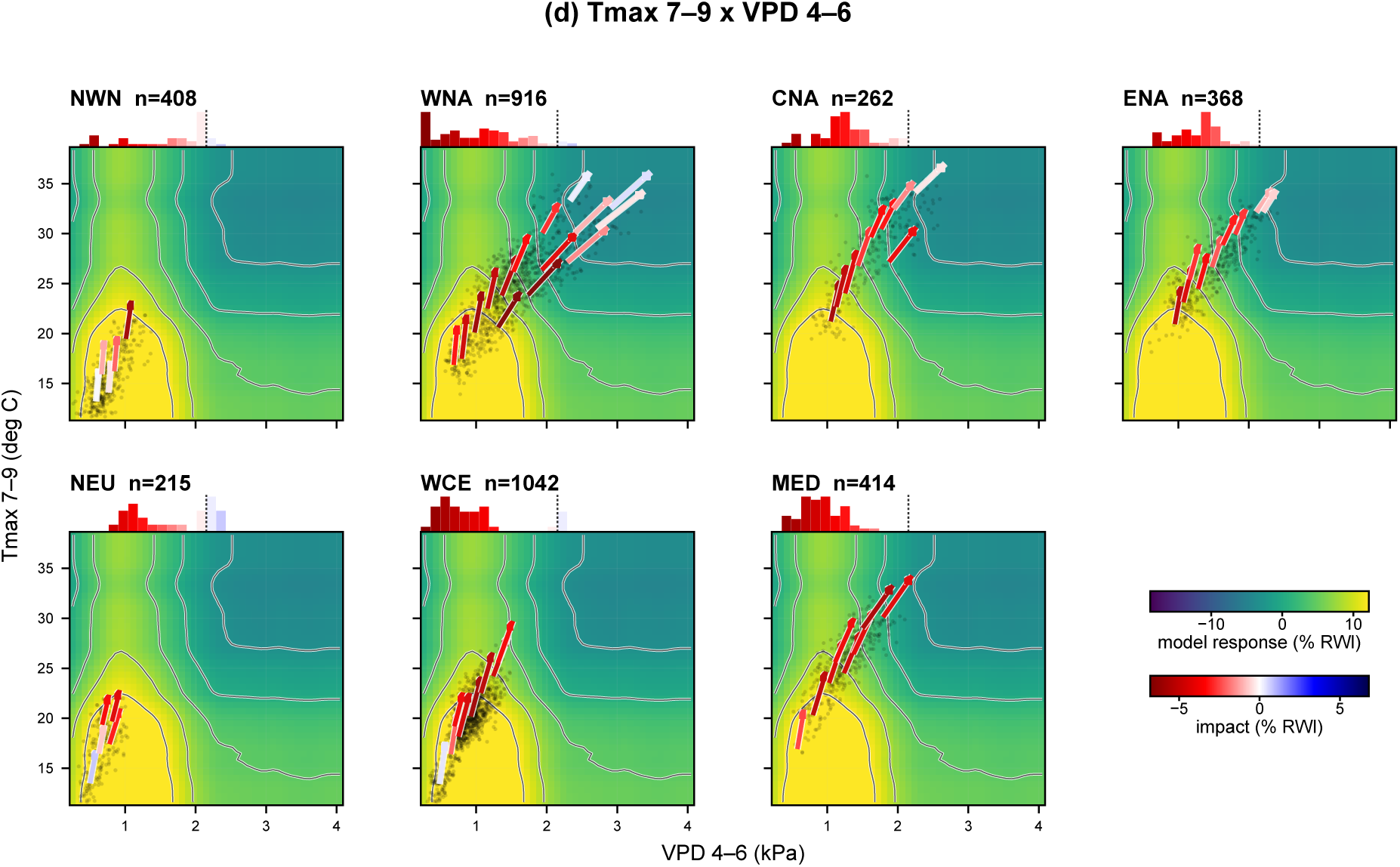
Regional response surfaces for D_4–6_ and T_7–9_. As Fig. S19, for the D_4–6_–T_7–9_ predictor pair.

**Figure S23:**
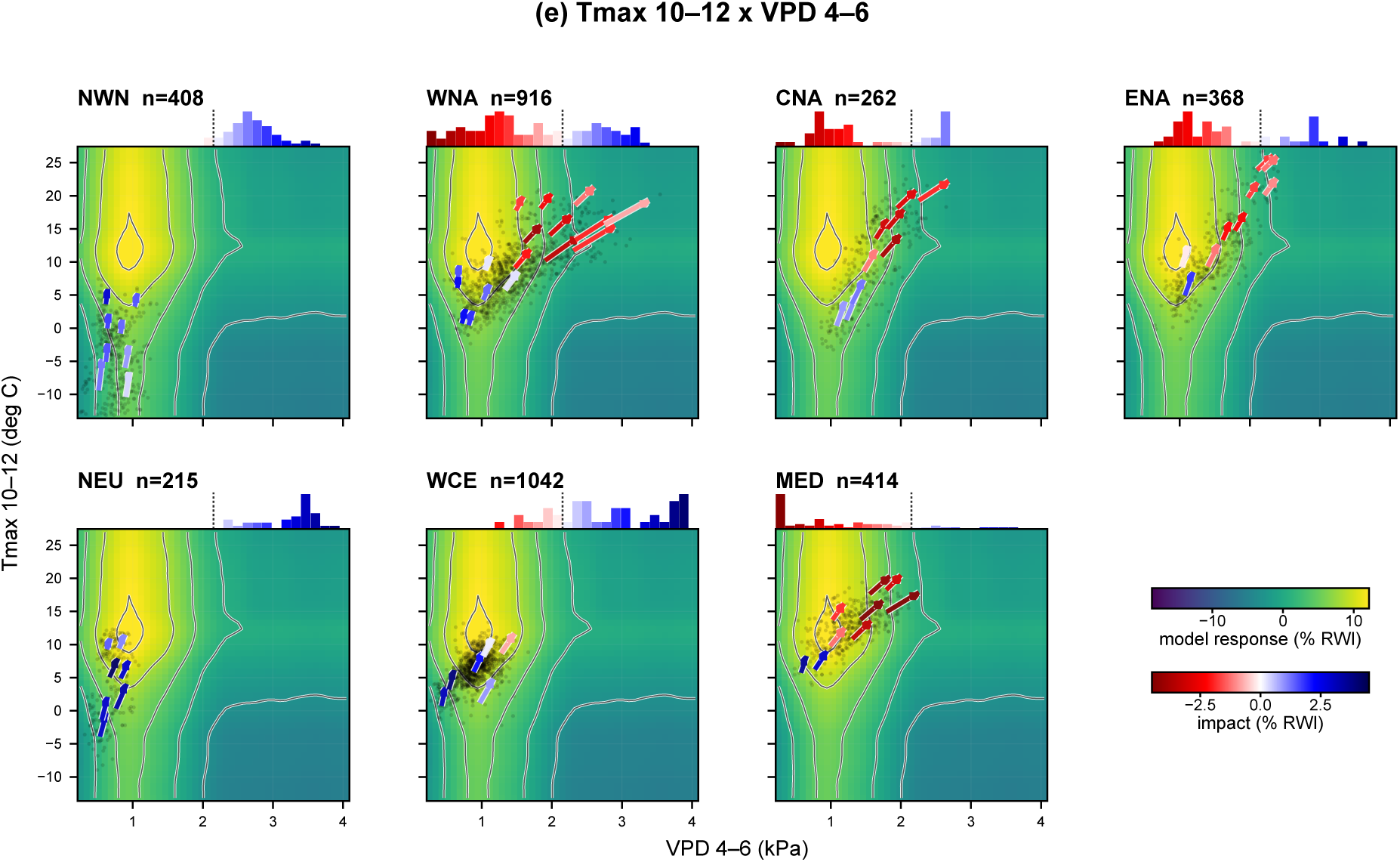
Regional response surfaces for D_4–6_ and T_10–12_. As Fig. S19, for the D_4–6_–T_10–12_ predictor pair.

**Figure S24:**
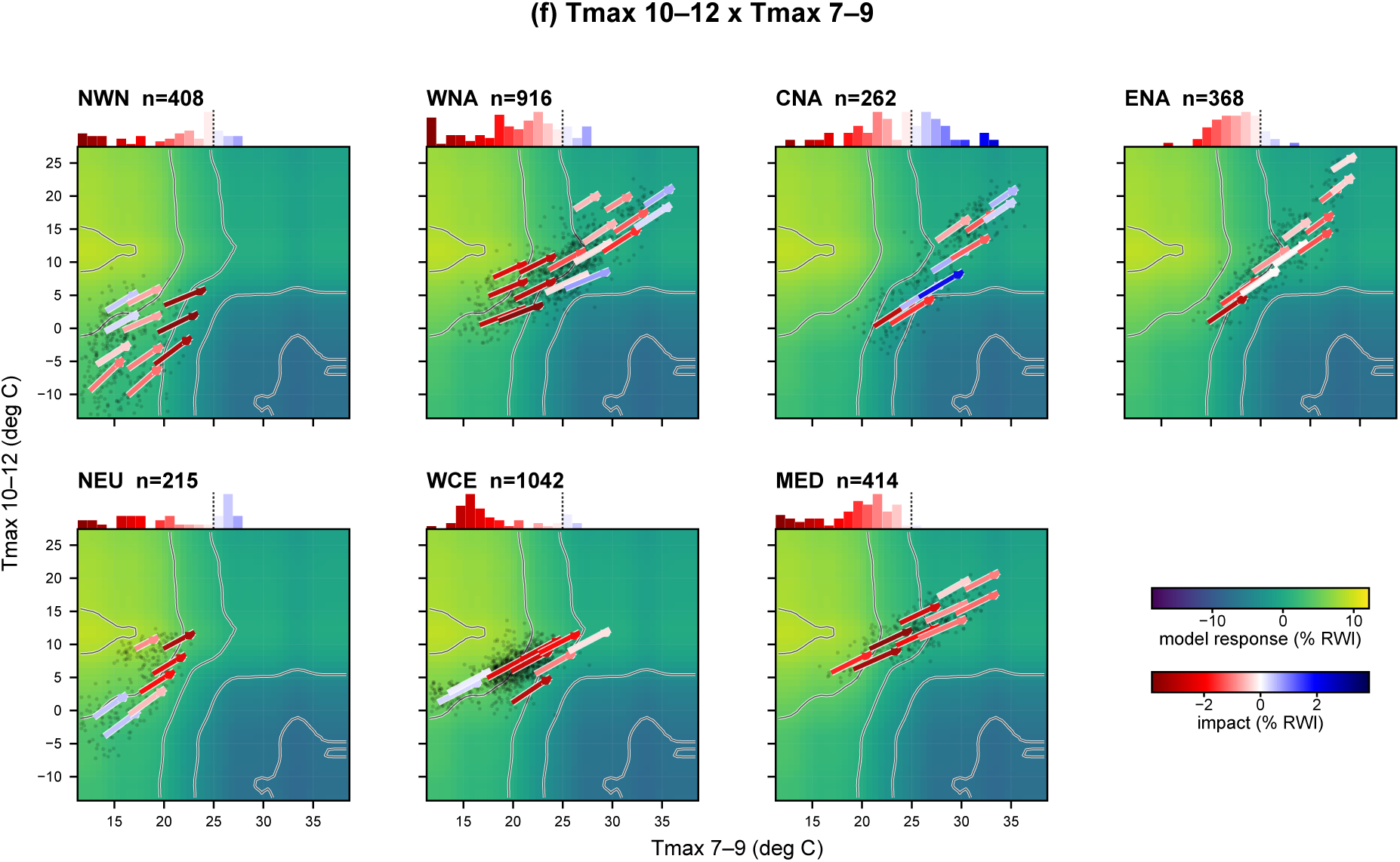
Regional response surfaces for T_7–9_ and T_10–12_. As Fig. S19, for the T_7–9_–T_10–12_ predictor pair.

## References

[1] William R. L. Anderegg, Anna T. Trugman, Grayson Badgley, Christa M. Anderson, Ann Bartuska, Philippe Ciais, Danny Cullenward, Christopher B. Field, Jeremy Freeman, Scott J. Goetz, Jeffrey A. Hicke, Deborah Huntzinger, Robert B. Jackson, Jordan Nickerson, Stephen Pacala, and James T. Randerson. Climate-driven risks to the climate mitigation potential of forests. Science, 368(6497):eaaz7005, 2020. doi: 10.1126/science.aaz7005.

[2] Flurin Babst, Olivier Bouriaud, Dario Papale, Bert Gielen, Ivan A Janssens, Eero Nikinmaa, Andreas Ibrom, Jian Wu, Christian Bernhofer, Barbara Köstner, et al. Above-ground woody carbon sequestration measured from tree rings is coherent with net ecosystem productivity at five eddy-covariance sites. New Phytologist, 201(4):1289–1303, 2014.

[3] Flurin Babst, Paul Bodesheim, Noah Charney, Andrew D. Friend, Martin P. Girardin, Stefan Klesse, David J. P. Moore, Kristina Seftigen, Jesper Björklund, Olivier Bouriaud, Andria Dawson, R. Justin DeRose, Michael C. Dietze, Annemarie H. Eckes, Brian Enquist, David C. Frank, Miguel D. Mahecha, Benjamin Poulter, Sydne Record, Valerie Trouet, Rachael H. Turton, Zhen Zhang, and Margaret E. K. Evans. When tree rings go global: Challenges and opportunities for retro- and prospective insight. Quaternary Science Reviews, 197:1–20, 2018. doi: 10.1016/j.quascirev.2018.07.009.

[4] Flurin Babst, Olivier Bouriaud, Benjamin Poulter, Valerie Trouet, Martin P. Girardin, and David C. Frank. Twentieth century redistribution in climatic drivers of global tree growth. Science Advances, 5(1):eaat4313, 2019. doi: 10.1126/sciadv.aat4313.

[5] Yoav Benjamini and Yosef Hochberg. Controlling the false discovery rate: a practical and powerful approach to multiple testing. Journal of the Royal statistical society: series B (Methodological), 57(1):289–300, 1995.

[6] Gordon B Bonan. Forests and climate change: forcings, feedbacks, and the climate benefits of forests. science, 320(5882):1444–1449, 2008.

[7] Roel J. W. Brienen, Lisa Caldwell, Louis Duchesne, Steven Voelker, Jonathan Barichivich, Marco Baliva, Gregório Ceccantini, Alfredo Di Filippo, Samuli Helama, Giuliano M. Locos-selli, Luciano Lopez, Gianluca Piovesan, Jochen Schöngart, Ricardo Villalba, and Emanuel Gloor. Forest carbon sink neutralized by pervasive growth-lifespan trade-offs. Nature Communications, 11:4241, 2020. doi: 10.1038/s41467-020-17966-z.

[8] Timothy J. Brodribb, Jennifer Powers, Hervé Cochard, and Brendan Choat. Hanging by a thread? forests and drought. Science, 368(6488):261–266, 2020.

[9] Antoine Cabon, Steven A Kannenberg, Altaf Arain, Flurin Babst, Dennis Baldocchi, Soumaya Belmecheri, Nicolas Delpierre, Rossella Guerrieri, Justin T Maxwell, Shawn McKenzie, et al. Cross-biome synthesis of source versus sink limits to tree growth. Science, 376(6594):758–761, 2022.

[10] Noah D Charney, Flurin Babst, Benjamin Poulter, Sydne Record, Valerie M Trouet, David Frank, Brian J Enquist, and Margaret EK Evans. Observed forest sensitivity to climate implies large changes in 21st century north american forest growth. Ecology letters, 19(9): 1119–1128, 2016.

[11] Tianqi Chen and Carlos Guestrin. Xgboost: A scalable tree boosting system. In Proceedings of the 22nd acm sigkdd international conference on knowledge discovery and data mining, pages 785–794, 2016.

[12] Edward R Cook. The decomposition of tree-ring series for environmental studies. Tree-Ring Bulletin, 1987.

[13] Edward R Cook and Kenneth Peters. The smoothing spline: a new approach to standardizing forest interior tree-ring width series for dendroclimatic studies. Tree-Ring Bulletin, 1981.

[14] L. D’orangeville, Louis Duchesne, Daniel Houle, Daniel Kneeshaw, Benoît Côté, and Neil Pederson. Northeastern north america as a potential refugium for boreal forests in a warming climate. Science, 352(6292):1452–1455, 2016.

[15] Loïc D’Orangeville, Daniel Houle, Louis Duchesne, Richard P. Phillips, Yves Bergeron, and Daniel Kneeshaw. Beneficial effects of climate warming on boreal tree growth may be transitory. Nature Communications, 9:3213, 2018. doi: 10.1038/s41467-018-05705-4.

[16] Bradley Efron. Bootstrap methods: another look at the jackknife. In Breakthroughs in statistics: Methodology and distribution, pages 569–593. Springer, 1992.

[17] Margaret EK Evans, Peter B Adler, Amy L Angert, Sharmila MN Dey, Martin P Girardin, Kelly A Heilman, Stefan Klesse, Daniel L Perret, Dov F Sax, Seema N Sheth, et al. Reconsidering space-for-time substitution in climate change ecology. Nature Climate Change, 15(8):809–812, 2025.

[18] Simone Fatichi, Sebastian Leuzinger, and Christian Körner. Moving beyond photosynthesis: from carbon source to sink-driven vegetation modeling. New Phytologist, 201(4):1086–1095, 2014.

[19] Simone Fatichi, Nadav Peleg, Theodoros Mastrotheodoros, Christoforos Pappas, and Gabriele Manoli. An ecohydrological journey of 4500 years reveals a stable but threatened precipitation–groundwater recharge relation around jerusalem. Science Advances, 7 (37):eabe6303, 2021.

[20] Charlotte Grossiord, Thomas N Buckley, Lucas A Cernusak, Kimberly A Novick, Benjamin Poulter, Rolf TW Siegwolf, John S Sperry, and Nate G McDowell. Plant responses to rising vapor pressure deficit. New phytologist, 226(6):1550–1566, 2020.

[21] Ian Harris, Timothy J Osborn, Phil Jones, and David Lister. Version 4 of the cru ts monthly high-resolution gridded multivariate climate dataset. Scientific data, 7(1):109, 2020.

[22] Zeke Hausfather and Glen P Peters. Emissions–the ‘business as usual’story is misleading. Nature, 577(7792):618–620, 2020.

[23] Sabrina Hempel, Katja Frieler, Lila Warszawski, Jacob Schewe, and Franziska Piontek. A trend-preserving bias correction–the isi-mip approach. Earth System Dynamics, 4(2): 219–236, 2013.

[24] Maialen Iturbide, José Manuel Gutiérrez, Lincoln Muniz Alves, Joaquín Bedia, Ezequiel Cimadevilla, Antonio S Cofiño, Ruth Cerezo-Mota, Alejandro Di Luca, Sergio Henrique Faria, Irina Gorodetskaya, et al. An update of ipcc climate reference regions for subcon-tinental analysis of climate model data: definition and aggregated datasets. Earth System Science Data Discussions, 2020:1–16, 2020.

[25] Wenzhe Jiao, Lixin Wang, William K Smith, Qing Chang, Honglang Wang, and Paolo D’Odorico. Observed increasing water constraint on vegetation growth over the last three decades. Nature Communications, 12(1):3777, 2021.

[26] Steven A. Kannenberg, Antoine Cabon, Flurin Babst, Soumaya Belmecheri, Nicolas Delpierre, Rossella Guerrieri, Justin T. Maxwell, Frederick C. Meinzer, David J. P. Moore, Christoforos Pappas, Masahito Ueyama, Danielle E. M. Ulrich, Steven L. Voelker, David R. Woodruff, and William R. L. Anderegg. Drought-induced decoupling between carbon uptake and tree growth impacts forest carbon turnover time. Agricultural and Forest Meteorology, 322:108996, 2022. doi: 10.1016/j.agrformet.2022.108996.

[27] Stefan Klesse, R Justin DeRose, Christopher H Guiterman, Ann M Lynch, Christopher D O’Connor, John D Shaw, and Margaret EK Evans. Sampling bias overestimates climate change impacts on forest growth in the southwestern united states. Nature communications, 9(1):5336, 2018.

[28] Stefan Klesse, Robert Justin DeRose, Flurin Babst, Bryan A Black, Leander DL Anderegg, Jodi Axelson, Ailene Ettinger, Hardy Griesbauer, Christopher H Guiterman, Grant Harley, et al. Continental-scale tree-ring-based projection of douglas-fir growth: Testing the limits of space-for-time substitution. Global Change Biology, 26(9):5146–5163, 2020.

[29] Stefan Klesse, Richard L Peters, Raquel Alfaro-Sánchez, Vincent Badeau, Claudia Bait-tinger, Giovanna Battipaglia, Didier Bert, Franco Biondi, Michal Bosela, Marius Budeanu, et al. No future growth enhancement expected at the northern edge for european beech due to continued water limitation. Global change biology, 30(10):e17546, 2024.

[30] Cameron C Lee and Matthew P Dannenberg. Frequencies of multivariate air masses drive tree growth. Journal of Geophysical Research: Biogeosciences, 128(3):e2022JG007064, 2023.

[31] Andrea H Lloyd, Paul A Duffy, and Daniel H Mann. Nonlinear responses of white spruce growth to climate variability in interior alaska. Canadian Journal of Forest Research, 43 (999):331–343, 2013.

[32] Scott M Lundberg and Su-In Lee. A unified approach to interpreting model predictions. Advances in neural information processing systems, 30, 2017.

[33] Scott M Lundberg, Gabriel Erion, Hugh Chen, Alex DeGrave, Jordan M Prutkin, Bala Nair, Ronit Katz, Jonathan Himmelfarb, Nisha Bansal, and Su-In Lee. From local explanations to global understanding with explainable ai for trees. Nature machine intelligence, 2(1): 56–67, 2020.

[34] Andreas Lundgren, Joachim Strengbom, Johannes Edvardsson, and Gustaf Granath. Unpacking climate effects on boreal tree growth: An analysis of tree-ring widths across temperature and soil moisture gradients. EGUsphere, 2025:1–27, 2025.

[35] Edurne Martinez del Castillo, Max CA Torbenson, Frederick Reinig, Ernesto Tejedor, Martín de Luis, and Jan Esper. Contrasting future growth of norway spruce and scots pine forests under warming climate. Global Change Biology, 30(11):e17580, 2024.

[36] Nate G McDowell, Craig D Allen, Kristina Anderson-Teixeira, Brian H Aukema, Ben Bond-Lamberty, Louise Chini, James S Clark, Michael Dietze, Charlotte Grossiord, Adam Hanbury-Brown, et al. Pervasive shifts in forest dynamics in a changing world. Science, 368(6494):eaaz9463, 2020.

[37] Hanna Meyer and Edzer Pebesma. Predicting into unknown space? estimating the area of applicability of spatial prediction models. Methods in Ecology and Evolution, 12(9): 1620–1633, 2021.

[38] Ariane Mirabel, Martin P. Girardin, Juha Metsaranta, Danielle Way, and Peter B. Reich. Increasing atmospheric dryness reduces boreal forest tree growth. Nature Communications, 14:6901, 2023. doi: 10.1038/s41467-023-42466-1.

[39] NOAA/WDS Paleoclimatology. NOAA/WDS Paleoclimatology - ITRDB Data Bank, 2014. URL https://www.ncei.noaa.gov/products/paleoclimatology/tree-ring. Accessed 2 August 2026.

[40] Kimberly A Novick, Darren L Ficklin, Paul C Stoy, Christopher A Williams, Gil Bohrer, A Christopher Oishi, Shirley A Papuga, Peter D Blanken, Asko Noormets, Benjamin N Sulman, et al. The increasing importance of atmospheric demand for ecosystem water and carbon fluxes. Nature climate change, 6(11):1023–1027, 2016.

[41] Kimberly A Novick, Darren L Ficklin, Charlotte Grossiord, Alexandra G Konings, Jordi Martínez-Vilalta, Walid Sadok, Anna T Trugman, A Park Williams, Alexandra J Wright, John T Abatzoglou, et al. The impacts of rising vapour pressure deficit in natural and managed ecosystems. Plant, Cell & Environment, 2024.

[42] OpenDendro. dplpy: Dendrochronology Program Library for Python, 2025. URL https://opendendro.org/python/. Documentation page dated 19 March 2025; accessed 2 August 2026.

[43] Yude Pan, Richard A Birdsey, Jingyun Fang, Richard Houghton, Pekka E Kauppi, Werner A Kurz, Oliver L Phillips, Anatoly Shvidenko, Simon L Lewis, Josep G Canadell, et al. A large and persistent carbon sink in the world’s forests. science, 333(6045):988–993, 2011.

[44] Drew MP Peltier, Jarrett J Barber, and Kiona Ogle. Quantifying antecedent climatic drivers of tree growth in the southwestern us. Journal of Ecology, 106(2):613–624, 2018.

[45] Hendrik Poorter, Karl J Niklas, Peter B Reich, Jacek Oleksyn, Pieter Poot, and Liesje Mommer. Biomass allocation to leaves, stems and roots: meta-analyses of interspecific variation and environmental control. New phytologist, 193(1):30–50, 2012.

[46] Rupert Seidl, Dominik Thom, Markus Kautz, Dario Martin-Benito, Mikko Peltoniemi, Giorgio Vacchiano, Jan Wild, Davide Ascoli, Michal Petr, Juha Honkaniemi, Manfred J. Lexer, Volodymyr Trotsiuk, Paola Mairota, Miroslav Svoboda, Marek Fabrika, Thomas A. Nagel, and Christopher P. O. Reyer. Forest disturbances under climate change. Nature Climate Change, 7:395–402, 2017. doi: 10.1038/nclimate3303.

[47] Bridget Thrasher, Weile Wang, Andrew Michaelis, and Ramakrishna Nemani. NEX-GDDP-CMIP6. Dataset, 2021.

[48] Bridget Thrasher, Weile Wang, Andrew Michaelis, Forrest Melton, Tsengdar Lee, and Ramakrishna Nemani. Nasa global daily downscaled projections, cmip6. Scientific data, 9 (1):262, 2022.

[49] Jan Tumajer, Jakub Kašpar, Jan Altman, Nela Altmanová, J Julio Camarero, Emil Cienciala, Vojtĕch Čada, Tomáš Čihák, Jiří Doležal, Pavel Fibich, et al. Longer growing seasons will not offset growth loss in drought-prone temperate forests of central-southeast europe. Nature Communications, 16(1):9535, 2025.

[50] Jiejie Wang, Anthony R Taylor, and Loïc D’Orangeville. Warming-induced tree growth may help offset increasing disturbance across the canadian boreal forest. Proceedings of the National Academy of Sciences, 120(2):e2212780120, 2023.

[51] Danielle A Way and RAM Oren. Differential responses to changes in growth temperature between trees from different functional groups and biomes: a review and synthesis of data. Tree physiology, 30(6):669–688, 2010.

[52] John W Williams, Stephen T Jackson, and John E Kutzbach. Projected distributions of novel and disappearing climates by 2100 ad. Proceedings of the National Academy of Sciences, 104(14):5738–5742, 2007.

[53] Martin Wilmking, Marieke Van Der Maaten-Theunissen, Ernst Van Der Maaten, Tobias Scharnweber, Allan Buras, Christine Biermann, Marina Gurskaya, Martin Hallinger, Jelena Lange, Rohan Shetti, et al. Global assessment of relationships between climate and tree growth. Global Change Biology, 26(6):3212–3220, 2020.

[54] Edwin B Wilson. Probable inference, the law of succession, and statistical inference. Journal of the American Statistical Association, 22(158):209–212, 1927.

[55] Wenping Yuan, Yi Zheng, Shilong Piao, Philippe Ciais, Danica Lombardozzi, Yingping Wang, Youngryel Ryu, Guixing Chen, Wenjie Dong, Zhongming Hu, Atul K. Jain, Chongya Jiang, Etsushi Kato, Shihua Li, Sebastian Lienert, Shuguang Liu, Julia E. M. S. Nabel, Zhangcai Qin, Timothy Quine, Stephen Sitch, William K. Smith, Fan Wang, Chaoyang Wu, Zhiqiang Xiao, and Song Yang. Increased atmospheric vapor pressure deficit reduces global vegetation growth. Science Advances, 5(8):eaax1396, 2019. doi: 10.1126/sciadv.aax1396.

[56] Pieter A Zuidema, Flurin Babst, Peter Groenendijk, Valerie Trouet, Abrham Abiyu, Rodolfo Acuña-Soto, Eduardo Adenesky-Filho, Raquel Alfaro-Sánchez, José Roberto Vieira Aragão, Gabriel Assis-Pereira, et al. Tropical tree growth driven by dry-season climate variability. Nature Geoscience, 15(4):269–276, 2022.

